# Perception of structurally distinct effectors by the integrated WRKY domain of a plant immune receptor

**DOI:** 10.1101/2021.07.28.454147

**Authors:** N Mukhi, H Brown, D Gorenkin, P Ding, AR Bentham, JDG Jones, MJ Banfield

## Abstract

Plants use intracellular immune receptors (NLRs) to detect pathogen-derived effector proteins. The Arabidopsis NLR pair RRS1-R/RPS4 confers disease resistance to different bacterial pathogens by perceiving structurally distinct effectors AvrRps4 from *Pseudomonas syringae* pv*. pisi* and PopP2 from *Ralstonia solanacearum* via an integrated WRKY domain in RRS1-R. How the WRKY domain of RRS1 (RRS1^WRKY^) perceives distinct classes of effector to initiate an immune response is unknown. We report here the crystal structure of the in planta processed C-terminal domain of AvrRps4 (AvrRps4^C^) in complex with RRS1^WRKY^. Perception of AvrRps4^C^ by RRS1^WRKY^ is mediated by the β2-β3 segment of RRS1^WRKY^ that binds an electronegative patch on the surface of AvrRps4^C^. Structure-based mutations that disrupt AvrRps4^C^/RRS1^WRKY^ interactions in vitro compromise RRS1/RPS4-dependent immune responses. We also show that AvrRps4^C^ can associate with the WRKY domain of the related but distinct RRS1B/RPS4B NLR pair, and the DNA binding domain of *At*WRKY41, with similar binding affinities. This work demonstrates how integrated domains in plant NLRs can directly bind structurally distinct effectors to initiate immunity.

**Significance:** This study reveals a mechanism of effector recognition by a plant NLR immune receptor that carries an integrated domain (ID) which mimics an authentic pathogen effector target. An Arabidopsis immune receptor carrying RRS1 and RPS4 NLR proteins detects the *Pseudomonas syringae* pv*. pisi* secreted effector AvrRps4 via a WRKY ID in RRS1. We used structural biology to reveal the mechanisms of AvrRps4/WRKY interaction and demonstrated that this binding is essential for effector recognition in planta. Our analysis revealed distinctive features of the WRKY ID that mediate the recognition of structurally distinct effectors from different bacterial pathogens. These insights could enable engineering NLRs with novel recognition specificities, and enhances our understanding of how effectors interact with host proteins.

## Introduction

Plants co-evolve with their pathogens, resulting in extensive genetic variation in host immune receptor and pathogen virulence factor (effector) repertoires (1). To enable host colonization, pathogenic microbes deliver effector proteins into host cells that suppress host immune responses and elevate host susceptibility by manipulating host physiology (2, 3). Plants have evolved surveillance mechanisms to detect and then activate defenses that combat pathogens, and detect host-translocated effectors via nucleotide-binding leucine-rich repeat receptors (NLRs) (4). NLR genes are highly diverse, showing both copy number and presence/absence variation, and different alleles can exhibit distinct pathogen effector recognition specificities (5, 6). Plant NLR alleles usually recognize a specific effector (as described by the gene-for-gene model (7)). However, NLRs capable of responding to multiple effectors are known (5, 8, 9).

NLRs typically contain an N-terminal Toll/Interleukin-1 receptor/Resistance (TIR) or coiled coil (CC or CC_R_) domain, a central nucleotide binding (NB-ARC) domain, and a C-terminal leucine-rich repeat (LRR) domain (6). In addition to these canonical domains, some NLRs have evolved to carry integrated domains that mimic effector virulence targets and facilitate immune activation by directly binding effectors (10–15). Interestingly, integrated domain-containing NLRs (NLR-IDs) usually function with a paired helper NLR, which is required for immune signaling (16, 17).

The Arabidopsis NLR pair RRS1-R/RPS4 is a particularly interesting NLR-ID/NLR pair that confers recognition-dependent resistance to bacterial pathogens *Pseudomonas syringae* and *Ralstonia solanacearum*, and also resistance to a fungal pathogen (*Colletotrichum higginsianum*) where the effector is unknown (18–21). RRS1-R contains an integrated WRKY domain near its C-terminus (RRS1^WRKY^), which interacts with two structurally distinct type III secreted bacterial effectors – AvrRps4 from *Pseudomonas syringae* pv*. pisi* and PopP2 from *Ralstonia solanacearum* (13, 14, 22, 23). The RRS1^WRKY^ domain may mimic WRKY transcription factors, the virulence-associated targets of AvrRps4 and PopP2 to detect the effectors for immune recognition (13). Two alleles of RRS1 have been identified that differ in the length of the C-terminal extension after the WRKY domain. RRS1-R from accession Ws-2 has a 101 amino acid C-terminal extension beyond the end of the WRKY domain, and can perceive AvrRps4 and PopP2, while RRS1-S from Col-0, which perceives AvrRps4 but not PopP2, is likely a derived allele with a premature stop codon, and has only an 18 amino acid C-terminal extension (24). Most Arabidopsis ecotypes also carry a paralogous and genetically linked RRS1B/RPS4B NLR pair, which only perceives AvrRps4 (25). RRS1B/RPS4B share a similar domain architecture with RRS1/RPS4, including 60% sequence identity in the integrated WRKY domain.

AvrRps4 is proteolytically processed in planta to produce a 133-amino-acid N-terminal fragment (AvrRps4^N^) and an 88-amino-acid C-terminal fragment (AvrRps4^C^) (26, 27). Previous studies have highlighted the role of AvrRps4^C^ in triggering RRS1/RPS4-dependent immune responses (26, 27). AvrRps4^N^ has been reported to potentiate immune signaling from AvrRps4^C^ (28, 29). PopP2 is sequence and structurally distinct from AvrRps4 and has an acetyltransferase activity that is likely related to its role in virulence. The structural basis of PopP2 perception by RRS1^WRKY^ has been determined (30), but how RRS1^WRKY^ binds AvrRps4^C^ and whether this is via a shared or different interface to PopP2, is unknown.

Here, we determined the structural basis of AvrRps4^C^ recognition by the RRS1/RPS4 immune pair. The recognition of AvrRps4^C^ is mediated by the β2-β3 segment of RRS1^WRKY^, the same region used to bind PopP2. This segment interacts with surface-exposed acidic residues of AvrRps4^C^. Structure-informed mutagenesis at the AvrRps4^C^/RRS1^WRKY^ interface identifies AvrRps4 residues required for protein/protein interactions in vitro and in planta, and AvrRps4 perception and immune responses. Residues mediating the interaction of AvrRps4^C^ and RRS1^WRKY^ are conserved in both the RRS1B^WRKY^ and the DNA binding domain of WRKY transcription factors, and AvrRps4^C^ mutants that prevent interaction with RRS1^WRKY^ also disrupt binding to *At*WRKY41. This supports the hypothesis that the RRS1^WRKY^ mimics host WRKY transcription factors via a shared effector binding mechanism.

## Results

### AvrRps4^C^ interacts with the integrated WRKY domain of RRS1 in vitro

To investigate how AvrRps4^C^ interacts with the RRS1^WRKY^ domain, constructs comprising residues 134-221 of AvrRps4^C^ (the in planta processed C-terminal fragment) and residues 1194-1273 of RRS1-R (corresponding to the RRS1^WRKY^ domain) were separately expressed in *E. coli* and proteins purified (*SI Materials and Methods*). We qualitatively assessed the interaction of purified AvrRps4^C^ with RRS1^WRKY^ using analytical gel filtration chromatography. Individually, the proteins displayed well-separated elution profiles. RRS1^WRKY^ eluted at a volume (V_e_) of 14.9 mL and AvrRps4^C^ at a V_e_ of 12.1 mL (Fig. 1A). Following incubation of a 1:1 molar ratio of the proteins we observe a new elution peak with an earlier V_e_ of 11.8 mL, and a lack of absorption peaks for the separate proteins (Fig. 1A). This demonstrates complex formation in vitro and suggests a 1:1 stoichiometry in the AvrRps4^C^/RRS1^WRKY^ complex.

**Fig. 1.**
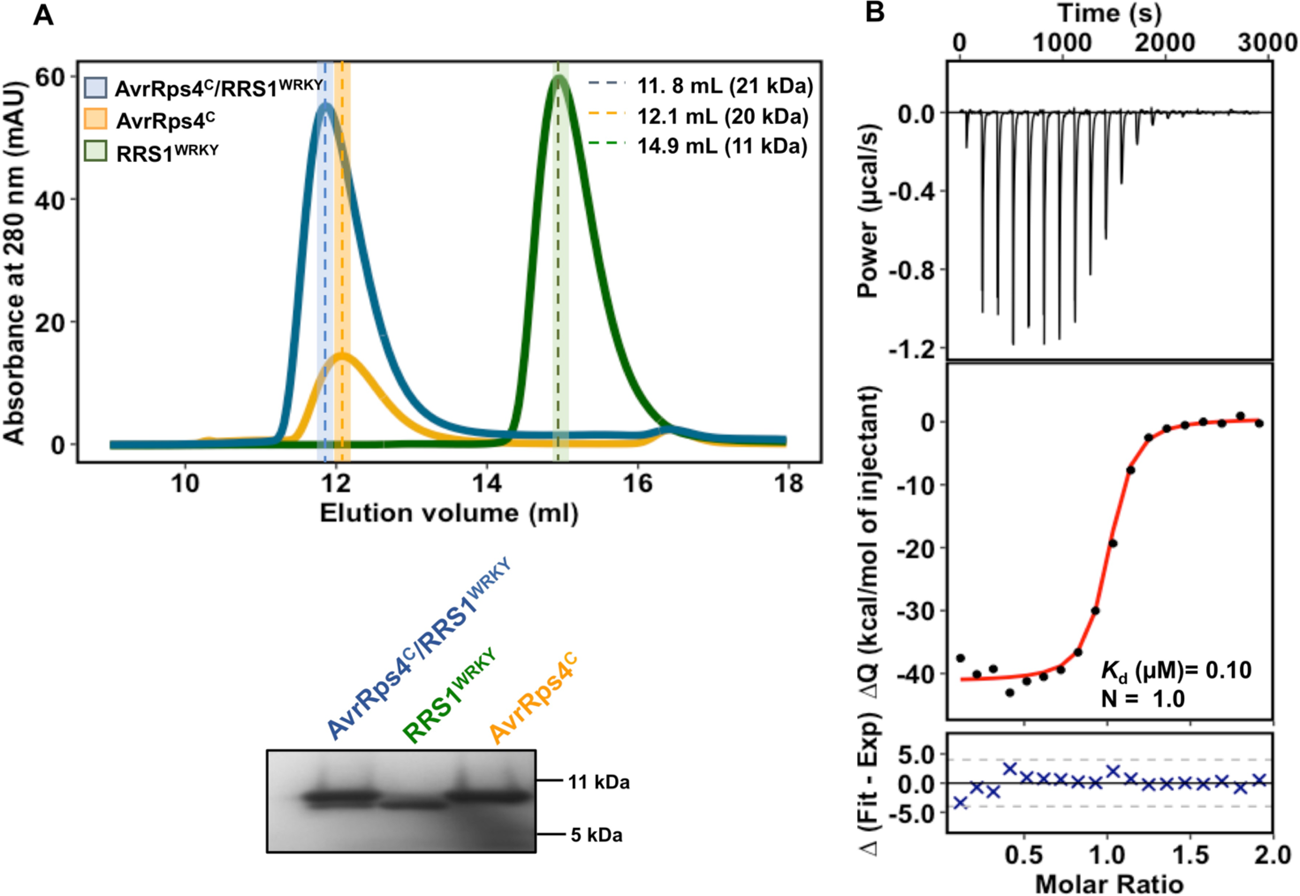
AvrRps4^C^ interacts with the WRKY domain of RRS1 in vitro. A) Analytical gel filtration traces for AvrRps4^C^ alone (Gold), RRS1^WRKY^ alone (green) and AvrRps4^C^ with RRS1^WRKY^ (Blue). An equimolar ratio of AvrRps4^C^ and RRS1^WRKY^ was used for the analysis. AvrRps4^C^ runs as a dimer *in vitro*. Poor absorbance for AvrRps4^C^ at 280nm is due its low molar extinction coefficient. (B) Isothermal titration calorimetry (ITC) titrations of AvrRps4^C^ with RRS1^WRKY^. Raw processed thermogram after baseline correction and noise removal is displayed in the upper panel. The lower panel represents the experimental binding isotherm obtained for the interaction of AvrRps4^C^ and RRS1^WRKY^ together with the global fitted curves (displayed in red) obtained from three independent experiments using Affinimeter software (60). *K*_d_ and binding stoichiometry (*N*) were derived from fitting to 1:1 binding model.

We then determined the binding affinities of the interaction using isothermal titration calorimetry (ITC). Titration of AvrRps4^C^ into a solution of RRS1^WRKY^ resulted in an exothermic binding isotherm with a fitted dissociation equilibrium constant (*K*_d_) of 0.103 μM (Fig. 1B) and stoichiometry of 1:1. The thermodynamic parameters of the interaction are given in Table 1. As RRS1^WRKY^ maybe a mimic of WRKY transcription factors, we explored the binding kinetics of AvrRps4^C^ with *At*WRKY41 and *At*WRKY70 by ITC (previous reports have shown that AvrRps4 interacts with these proteins in yeast two-hybrid and by in planta co-immunoprecipitation (13, 31)). We chose *At*WRKY41 for further study as this protein expressed and purified stably from *E. coli*. AvrRps4^C^ interacted with *At*WRKY41 with a *K*_d_ of 0.02 μM, and similar thermodynamic parameters as RRS1^WRKY^ (Fig. S1, Table 1).

**Table 1.** Thermodynamic parameters obtained from ITC experiments

| Cell | | Syringe | | T | $\Delta H$ | $K_d$ |
| --- | --- | --- | --- | --- | --- | --- |
| Contents | Conc.<br>[mM] | Contents | Conc.<br>[mM] | [°C] | [kcal mol<sup>-1</sup>] | [M<sup>-6</sup>] |
| RRS1<sup>WRKY</sup> | 0.02 | AvrRps4<sup>C</sup> | 0.2 | 25 | -40 ± 0.036 | 0.10 |
| RRS1<sup>WRKY</sup> | 0.02 | D164A | 0.2 | 25 | n.b. | n.b. |
| RRS1<sup>WRKY</sup> | 0.02 | N171A | 0.2 | 25 | -25.42 ± 0.02 | 0.04 |
| RRS1<sup>WRKY</sup> | 0.02 | E175A | 0.2 | 25 | n.b. | n.b. |
| RRS1<sup>WRKY</sup> | 0.02 | E187A | 0.2 | 25 | n.b. | n.b. |
| RRS1<sup>WRKY</sup> | 0.02 | Q194A | 0.2 | 25 | -19.89 ± 0.25 | 0.70 |
| RRS1<sup>WRKY</sup> | 0.02 | EE/AA | 0.2 | 25 | n.b. | n.b. |
| RRS1<sup>WRKY</sup> | 0.02 | KRVY/AAAA | 0.2 | 25 | -19.59 ± 0.02 | 0.03 |
| RRS1B<sup>WRKY</sup> | 0.02 | AvrRps4<sup>C</sup> | 0.2 | 25 | -14.48 ± 0.01 | 0.33 |
| WRKY41 | 0.02 | AvrRps4<sup>C</sup> | 0.2 | 25 | -90.54 ± 0.44 | 0.02 |
| WRKY41 | 0.02 | AvrRps4EE/AA | 0.2 | 25 | n.b. | n.b. |

### Crystal structure of the AvrRps4^C^/RRS1^WRKY^ complex

To reveal the molecular basis of AvrRps4^C^ and RRS1^WRKY^ interaction, we co-expressed the proteins in *E. coli*, purified the complex and obtained crystals that diffracted to 2.65 Å resolution at the Diamond Light Source, UK (see *SI Materials and Methods*). The crystal structure of the AvrRps4^C^/RRS1^WRKY^ complex was solved by molecular replacement using the structure of RRS1^WRKY^ (from the PopP2/RRS1^WRKY^ complex PDB ID: 5W3X) and AvrRps4^C^ (PDB ID: 4B6X) as models (see *SI Materials and Methods*). X-ray data collection, refinement and validation statistics are shown in Table 2.

**Table 2.** Data collection and refinement statistics for the crystal structure of the AvrRps4^C^/RRS1^WRKY^ complex.

| **Data collection statistics** |  |
| --- | --- |
| Wavelength (Å) | 0.912 |
| Space group | *P* 6<sub>1</sub> 2 2 |
| Cell dimensions a, b, c (Å) | 105.61, 105.61, 66.99 |
| Resolution (Å)* | 45.77 - 2.65 (2.78 - 2.65) |
| R<sub>merge</sub> (%) | 9.5 (119.0) |
| (I)/σ(I) | 17.4 (2.3) |
| Completeness (%) | 99.9 (100) |
| Unique reflections | 6800 (888) |
| Redundancy | 20.8 (21.9) |
| CC<sup>1/2</sup> | 1.0 (0.9) |
| **Refinement and model statistics** |  |
| Resolution (Å)* | 45.77 - 2.65 (2.72 - 2.65) |
| *R*<sub>work</sub> / *R*<sub>free</sub> (%) | 24.5 (35.3) / 28.3 (45.1) |
| No. atoms | 1020 |
| Protein | 1016 |
| Water | 3 |
| Mean B value (overall Å<sup>2</sup>) | 93.11 |
| rmsd bond lengths (Å) | 0.0039 |
| rmsd bond angles (°) | 1.24 |
| Ramachandran plot (%)** |  |
| Favoured | 95.9% |
| Allowed | 4.1% |
| Outliers | 0 |
| MolProbity Score | 1.89 (98<sup>th</sup> percentile) |

The structure comprises a 1:1 complex of AvrRps4^C^ and RRS1^WRKY^ (Fig. 2A), consistent with the stoichiometry suggested by analytical gel filtration and from ITC. Overall, AvrRps4^C^ adopts the same antiparallel α-helical coiled coil structure in both free (PDB ID: 4B6X (27)) and complexed forms, with an RMSD of 0.66 Å over 59 C*_α_* atoms (Fig S2A). Also, RRS1^WRKY^ adopts a conventional WRKY domain fold (RMSD of 2.03 Å over 61 C*_α_* atoms compared to *At*WRKY1, PDB ID: 2AYD (32)) comprising a four-stranded antiparallel β-sheet (β2 to β5) stabilized by a zinc ion (C_2_H_2_ type). Comparison of RRS1^WRKY^ in the AvrRps4^C^/RRS1^WRKY^ and PopP2/RRS1^WRKY^ complex (PDB ID: 5W3X) structures reveals high conformational similarity, with an RMSD of 1.81 Å over 64 C*_α_* atoms. The characteristic WRKY sequence signature motif ‘WRKYGQK’ maps to the β2 strand of RRS1^WRKY^ and is directly involved in contacting AvrRps4^C^ (Fig. 2B, Fig. S2B). The same surface, including the β2-β3 strands of RRS1^WRKY^, forms contacts with PopP2 in the structure of the PopP2/RRS1^WRKY^ complex (30) (Fig. S3) and mutants at this surface showed it to be essential for PopP2 recognition.

**Fig. 2.**
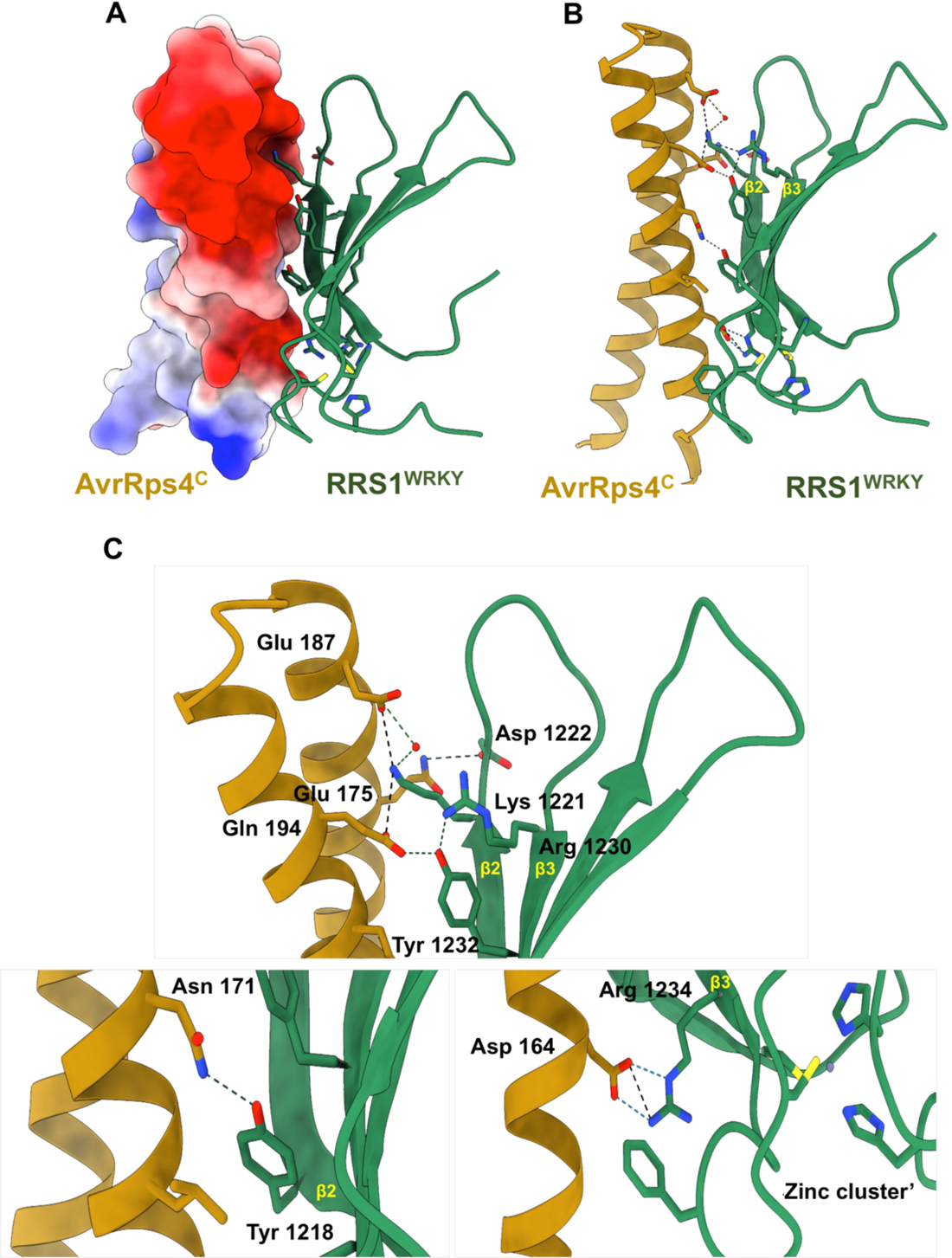
Structure of the AvrRps4^C^/RRS1^WRKY^ complex. (A) Electrostatic surface representation of AvrRps4^C^ in AvrRps4^C^/RRS1^WRKY^ crystal structure displaying prominent negative patch in AvrRps4 at the interacting interface. (B) Schematic representation of AvrRps4^C^/RRS1^WRKY^, highlighting interfacing residues. AvrRps4^C^ is shown in gold cartoon and RRS1^WRKY^ is shown in green with surface exposed side chains as sticks. (C) Close-up view of the interactions of AvrRps4^C^ with β2, β3 segment of RRS1^WRKY^. Hydrogen bonds are shown as dashed lines, and water molecules depicted as red spheres. The Zn^2+^ ion is also displayed.

### The AvrRps4^C^/RRS1^WRKY^ binding interface is dominated by electrostatic and polar interactions

The total interface area buried in the AvrRps4^C^/RRS1^WRKY^ complex is 591.8 Å^2^, encompassing 12.3 % (589.7 Å^2^) and 11.9 % (593.9 Å^2^) of the total accessible surface areas of the effector and integrated domain respectively (as calculated by PDBePISA (33), full details are given in Table 3). The binding interface between AvrRps4^C^ and RRS1^WRKY^ is largely formed by residues from the β2-β3 strand of RRS1^WRKY^, which present a positive surface patch that interacts with acidic residues on the surface of AvrRps4^C^ (Fig. 2A, Fig. S2B). The interaction between the β2 segment of RRS1^WRKY^, which harbors the WRK^1^YGQK^2^ motif, and AvrRps4^C^, includes hydrogen bond and/or salt bridge interactions involving Tyr1218 and Lys(K^2^)1221 of RRS1^WRKY^ and AvrRps4 Glu175, Glu187 and Asn171. Notably, the side chain of RRS1^WRKY^ Lys1221 protrudes into an acidic cleft on the surface of AvrRps4^C^ to contact the side chains of both AvrRps4 Glu175 and Glu187 (Fig. 2B,C). The OH atom of RRS1^WRKY^ Tyr1218 forms a hydrogen bond with the ND2 atom of AvrRps4 Asn171 (Fig. 2B,C). Additional intermolecular contacts are formed by the β2-β3 loop of RRS1^WRKY^ involving the backbone carbonyl oxygen and nitrogen of Asp1222, which form hydrogen bonds with the side chains of AvrRps4 Asn190 and Gln194. The complex between AvrRps4^C^/RRS1^WRKY^ is further stabilized by the β3 strand of RRS1^WRKY^ that forms hydrogen bonds and salt bridge interactions via side chains of RRS1^WRKY^ Arg1230, Tyr1232, Arg1234 to AvrRps4 Glu175 and Asp164 (Fig. 2B,C). A detailed interaction summary is provided in Table 4.

**Table 3.**
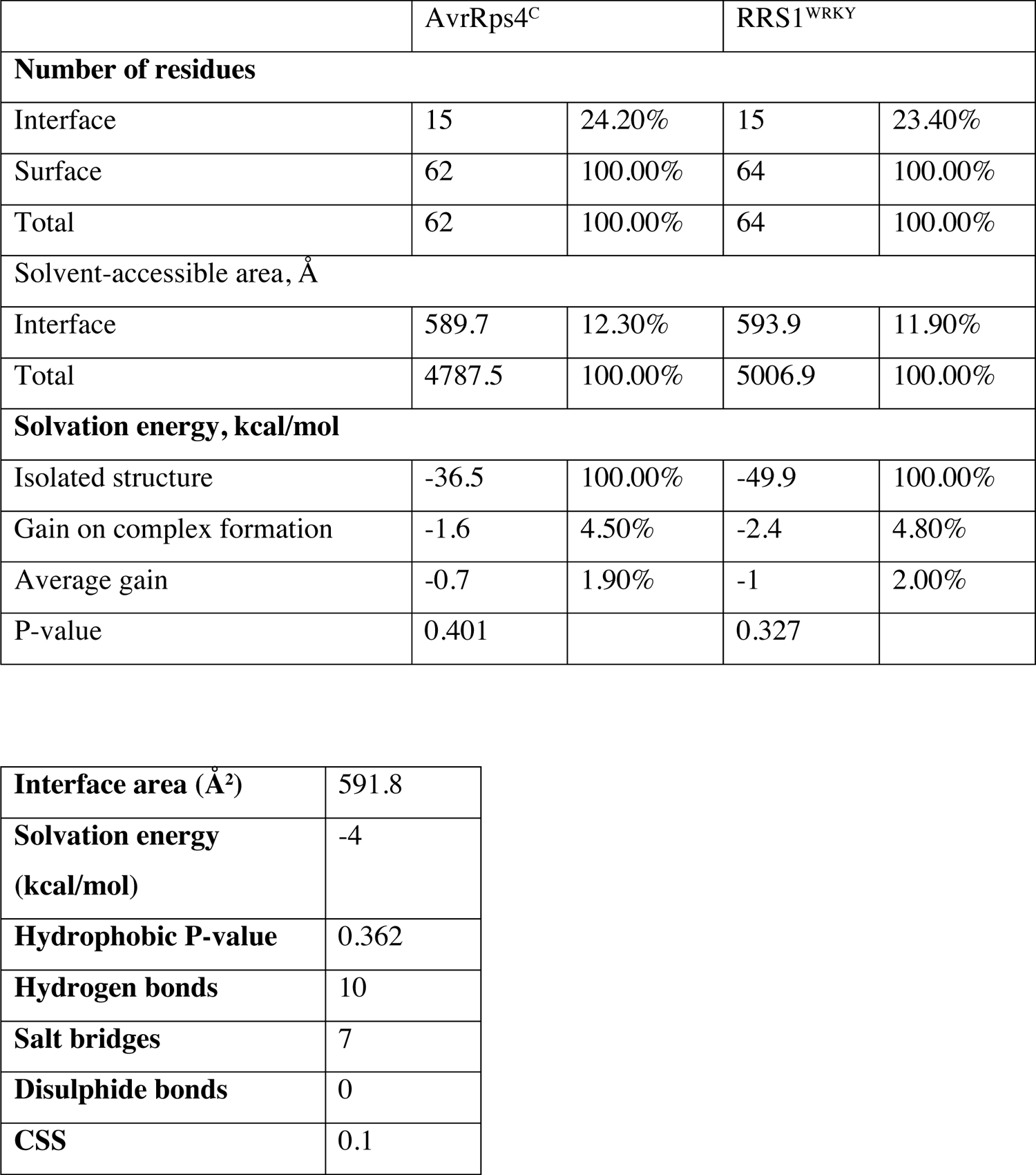
Interface summary for AvrRps4^C^/RRS1^WRKY^ complex. Interface analysis was performed using PDBePISA

|  |  |  |  |  |
| --- | --- | --- | --- | --- |
|  | AvrRps4<sup>C</sup> |  | RRS1<sup>WRKY</sup> |  |
| Number of residues |  |  |  |  |
| Interface | 15 | 24.20% | 15 | 23.40% |
| Surface | 62 | 100.00% | 64 | 100.00% |
| Total | 62 | 100.00% | 64 | 100.00% |
| Solvent-accessible area, Å |  |  |  |  |
| Interface | 589.7 | 12.30% | 593.9 | 11.90% |
| Total | 4787.5 | 100.00% | 5006.9 | 100.00% |
| Solvation energy, kcal/mol |  |  |  |  |
| Isolated structure | -36.5 | 100.00% | -49.9 | 100.00% |
| Gain on complex formation | -1.6 | 4.50% | -2.4 | 4.80% |
| Average gain | -0.7 | 1.90% | -1 | 2.00% |
| P-value | 0.401 |  | 0.327 |  |

|  |  |
| --- | --- |
| **Interface area (Å<sup>2</sup>)** | 591.8 |
| **Solvation energy (kcal/mol)** | -4 |
| **Hydrophobic P-value** | 0.362 |
| **Hydrogen bonds** | 10 |
| **Salt bridges** | 7 |
| **Disulphide bonds** | 0 |
| **CSS** | 0.1 |

**Table 4.** Interaction summary of the residues mediating the intermolecular contacts between AvrRps4^C^ and RRS1^WRKY^.

| **Hydrogen bonds** |  |  |  |
| --- | --- | --- | --- |
| **AvrRps4<sup>C</sup>** |  | **Dist. [Å]** | **RRS1<sup>WRKY</sup>** |
| 1 | A:ASN 171[ ND2] | 2.68 | B:TYR1218[ OH ] |
| 2 | A:ASN 190[ ND2] | 2.93 | B:ASP1222[ O ] |
| 3 | A:GLN 194[ NE2] | 2.99 | B:ASP1222[ O ] |
| 4 | A:GLU 187[ OE2] | 3.32 | B:LYS1221[ NZ ] |
| 5 | A:THR 191[ OG1] | 3.00 | B:LYS1221[ NZ ] |
| 6 | A:GLN 194[ OE1] | 2.90 | B:ASP1222[ N ] |
| 7 | A:GLU 175[ OE2] | 3.24 | B:ARG1230[ NH2] |
| 8 | A:GLU 175[ OE2] | 2.63 | B:TYR1232[ OH ] |
| 9 | A:ASP 164[ OD1] | 2.71 | B:ARG1234[ NE ] |
| 10 | A:ASP 164[ OD2] | 2.61 | B:ARG1234[ NH2] |
| **Salt bridges** |  |  |  |
| **AvrRps4<sup>C</sup>** |  | **Dist. [Å]** | **RRS1<sup>WRKY</sup>** |
| 1 | A:GLU 187[ OE2] | 3.32 | B:LYS1221[ NZ ] |
| 2 | A:GLU 175[ OE1] | 3.50 | B:LYS1221[ NZ ] |
| 3 | A:GLU 175[ OE2] | 3.24 | B:ARG1230[ NH2] |
| 4 | A:ASP 164[ OD1] | 2.71 | B:ARG1234[ NE ] |
| 5 | A:ASP 164[ OD2] | 3.45 | B:ARG1234[ NE ] |
| 6 | A:ASP 164[ OD1] | 3.41 | B:ARG1234[ NH2] |
| 7 | A:ASP 164[ OD2] | 2.61 | B:ARG1234[ NH2] |

### Structure-based mutations in AvrRps4^C^ perturb binding to RRS1^WRKY^ in vitro

To evaluate the contribution of residues at the AvrRps4^C^/RRS1^WRKY^ interface to complex formation in vitro, we generated six structure-guided mutants in AvrRps4^C^ (native amino acid to Ala) and tested the effect on protein interactions by ITC. Each AvrRps4^C^ mutant was purified from *E. coli* under the same conditions as for the wild-type protein, and proper folding evaluated by circular dichroism (CD) spectroscopy (Fig. S4). ITC titrations were carried out as for the wild-type interactions. Individual ITC isotherms are shown in Fig. 3, and the thermodynamic parameters of the interactions are shown in Table 1. We found that mutating AvrRps4 residues Asp164, Glu175, Glu187, and double mutant Glu175/Glu187, essentially abolished complex formation in vitro (Fig. 3). Mutations in residues Asn171 and Gln194 retained binding to RRS1^WRKY^, with Asn171Ala displaying wild-type levels and Gln194Ala showing an ∼7-fold reduction in affinity. Besides structure-guided mutants, we also tested binding of an AvrRps4 KRVY/AAAA mutant, carrying mutations in the N-terminal KRVY motif (26), with RRS1^WRKY^. Unlike most interface mutants, AvrRps4 KRVY/AAAA retained wild-type-like binding affinity with RRS1^WRKY^ (Fig. 3).

**Fig. 3.**
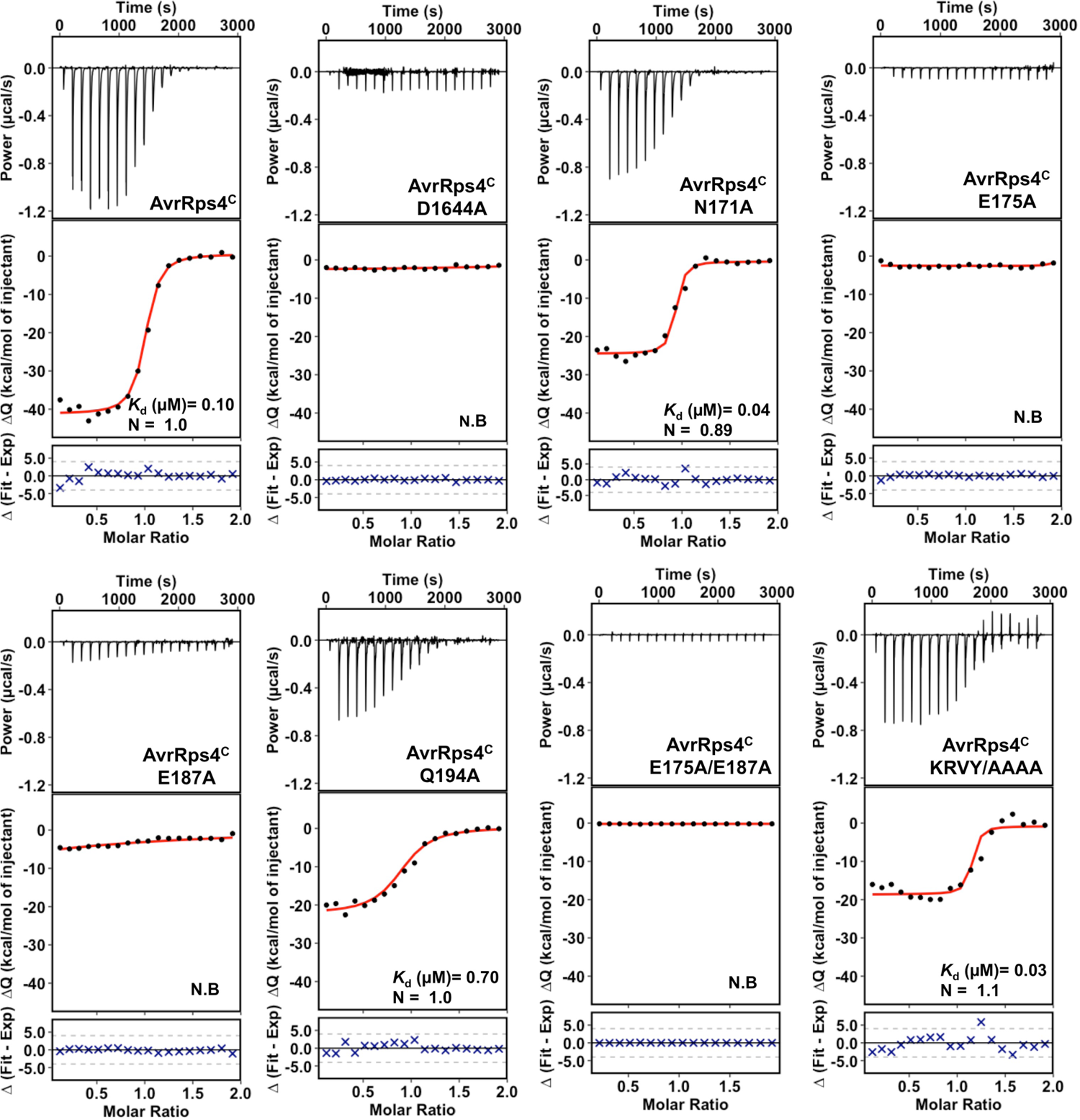
Structure-guided mutants of AvrRps4^C^ at the AvrRps4^C^/RRS1^WRKY^ interface disrupts interaction with the RRS1^WRKY^ in vitro. Isothermal titration calorimetry (ITC) titrations of wild-type AvrRps4^C^ and mutants with RRS1^WRKY^. Upper panels: raw processed thermograms after baseline correction and noise removal. Lower panels: experimental binding isotherm obtained for the interaction of AvrRps4^C^ wild-type and mutants with RRS1^WRKY^ together with the global fitted curves (displayed in red) obtained from three independent experiments using Affinimeter software (60). *K*_d_ was derived from fitting to 1:1 binding model.

Since AvrRps4^C^ associates with RRS1^WRKY^ and *At*WRKY41 with similar binding affinities (Fig. S1), we tested the impact of the AvrRps4 Glu175/Glu187 double mutant on the binding to *At*WRKY41. We found that this mutant also abolishes interaction with *At*WRKY41, suggesting the same AvrRps4 binding interface is shared with different WRKY proteins (Fig. S1).

### Structure-based mutations in AvrRps4 prevent RRS1/RPS4 mediated cell death in ***Nicotiana tabacum***

To validate the biological relevance of the AvrRps4^C^/RRS1^WRKY^ interface observed in the crystal structure, we tested the effect of the AvrRps4^C^ interface mutants above on RRS1-R/RPS4 mediated immunity by monitoring the cell-death response in *N. tabacum*. *Agrobacterium*-mediated transient expression of wild-type AvrRps4 triggers a hypersensitive cell death response (HR) 5 days post infiltration (dpi) when co-expressed with RRS1-R/RPS4 (Fig. 4A). The previously characterized inactive AvrRps4 KRVY/AAAA mutant (26, 27) was used as a negative control. We found that AvrRps4 mutations at positions Asp164, Glu175 and Glu187, and the double mutant Glu175/Glu187, prevented RRS1-R/RPS4-dependent cell death responses (Fig. 4A). Interestingly, the Asn171Ala mutation displayed wild-type like cell death-inducing activity, and Gln194Ala consistently exhibited weaker death. Expression of all mutants was confirmed by immunoblotting (Fig. 4B). In addition to RRS1-R/RPS4, we also explored the effect of AvrRps4 structure-based mutations on RRS1-S/RPS4-dependent cell death in *N. tabacum* (Fig. S5A). We found that AvrRps4 variants elicited similar immune responses when transiently co-expressed with RRS1-S/RPS4 or RRS1-R/RPS4.

**Fig. 4.**
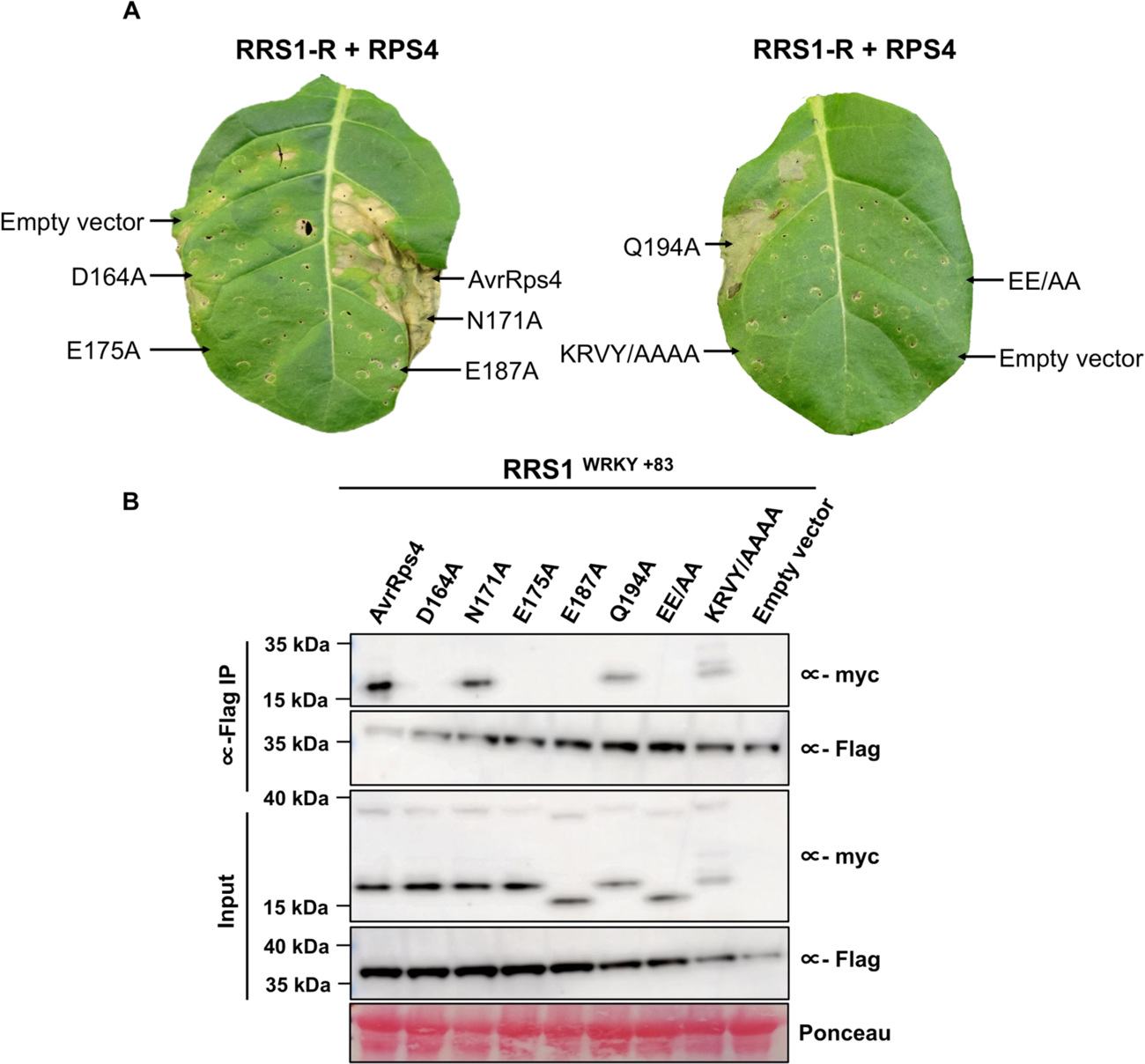
Structure-guided mutants of AvrRps4 at the AvrRps4^C^/RRS1^WRKY^ interface compromises RRS1-R/RPS4 mediated cell death responses and in vivo binding in *Nicotiana*. (A) Representative leaf images showing RRS1-R/RPS4 mediated cell death response to wild-type structure-guided mutants of AvrRps4. Agroinfiltration assays were performed in 4- to 5-week-old *N. tabacum* leaves, and cell death was assessed at 4 dpi. The experiment was repeated three times with similar results. (B) Co-IP of RRS1-R^WRKY+83^ (6xHis/3xFLAG-tagged) with AvrRps4^C^ and variants (4xmyc-tagged) in *N. benthamiana*. Blots show protein accumulations in total protein extracts (input) and immunoprecipitates obtained with anti-FLAG magnetic beads when probed with appropriate antisera. Empty vector was used as a control. The experiment was repeated at least three times, with similar results.

### Loss of cell death in *N. tabacum* correlates with the loss of binding to RRS1^WRKY^ in vivo

To determine whether loss of RRS1-R/RPS4-mediated HR in transient assays correlates with the loss of AvrRps4 binding to RRS1^WRKY^ in vivo, we performed co-immunoprecipitation (co-IP) assays using full-length C-terminal 4xmyc tagged AvrRps4 constructs and C-terminal 6xHis/3xFLAG-tagged constructs of RRS1-R^WRKY+83^ (RRS1-R^WRKY^ with an additional 83 amino acid at the C-terminal end, which enhances the stability of RRS1^WRKY^ when expressed in planta). Wild-type AvrRps4 associates with RRS1-R^WRKY+83^ in its in planta processed form (Fig. 4B). Consistent with the cell death phenotype, no association between AvrRps4 mutants Asp164Ala (D164A), Glu175Ala (E175A), Glu187Ala (E187A) or Glu175/187Ala (EE/AA) and RRS1^WRKY+83^ was detected (Fig. 4B). Further, we observed wild-type levels of association of AvrRps4 Asn171Ala (N171A) with RRS1^WRKY+83^, while AvrRps4 Gln194Ala (Q194A) appears to only co-IP weakly. The AvrRps4 KRVY/AAAA mutant displayed wild-type like binding affinity towards RRS1^WRKY+83^, as observed previously (27).

### Structure-guided mutations in AvrRps4 prevent HR in *A. thaliana*

Next, we investigated the impact of AvrRps4 structure-guided mutations on the activation of RRS1-R/RPS4-dependent immune responses using HR assays in *A. thaliana*. Constructs carrying full-length AvrRps4 wild-type and mutants, flanked by 126 bp native AvrRps4 promoter, were delivered into plant cells by infiltration using the Pf0-EtHAn (*Pseudomonas fluorescens* Effector-to-Host Analyzer, hence Pf0) system (34). HR assays used Arabidopsis ecotype Ws-2 (encoding RRS1-R/RPS4 and RPS4B/RRS1B) and Ws-2 *rrs1-1/rps4-21/rps4b-1* (RRS1-R/RPS4/RPS4B triple knockout) lines and scored at 20 hpi (hours post infiltration). Pf0 carrying wild-type AvrRps4 triggered HR in Ws-2, but not in Ws-2 *rrs1-1/rps4-21/rps4b-1,* as previously reported (13, 27). AvrRps4 KRVY/AAAA, an HR inactive mutant, was used as a negative control (27). The structure-guided mutants AvrRps4 D164A, E175A, E187A and EE/AA all showed a complete loss of HR in Ws-2, with AvrRps4 Q194A showing a weaker HR and N171A a wild-type-like phenotype (Fig. 5A). None of the AvrRps4 variants triggered HR in Ws-2 *rrs1-1/rps4-21/rps4b-1* (Fig. 5A).

**Fig. 5.**
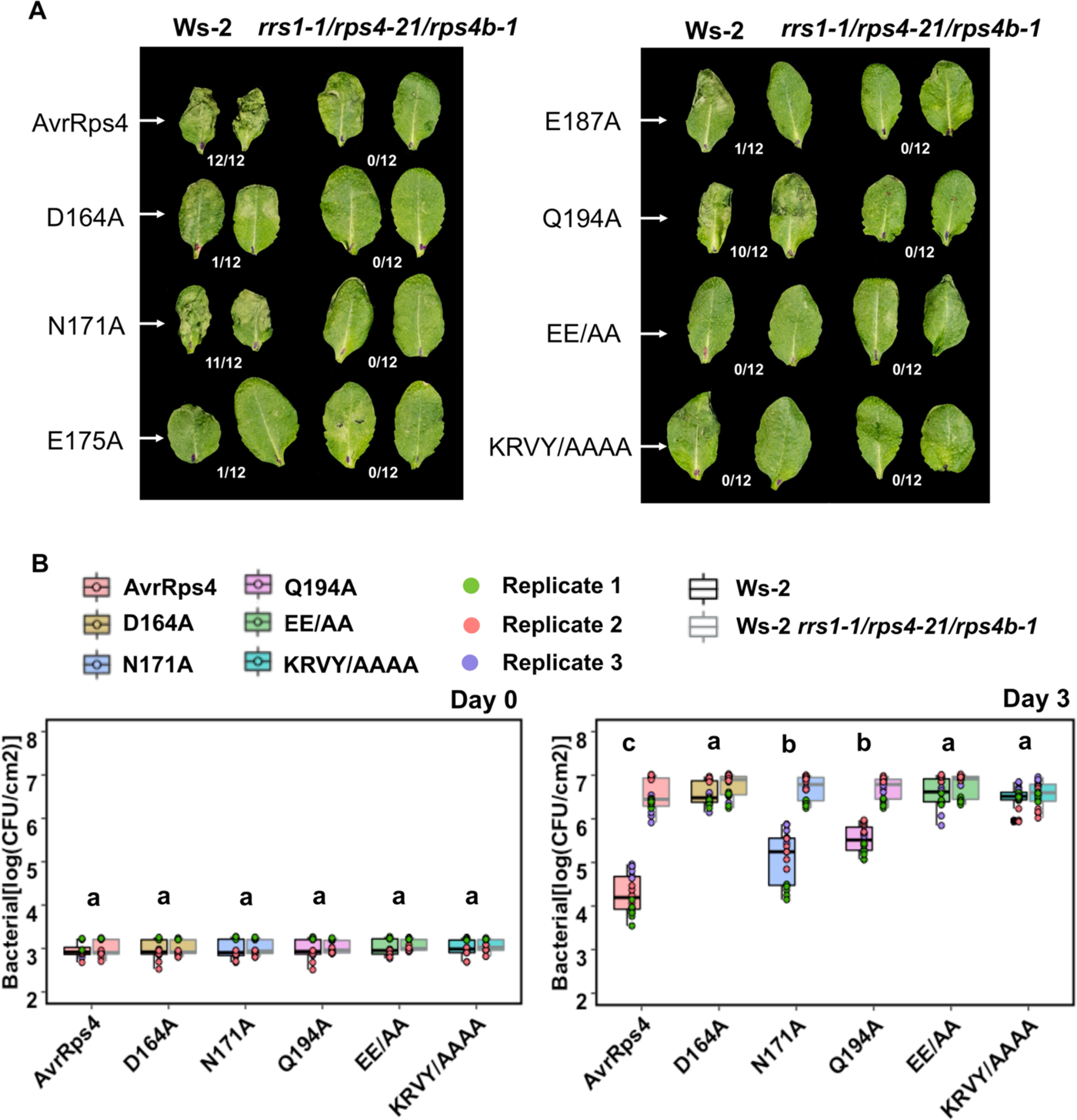
Structural-guided mutants of AvrRps4 compromises RRS1-R/RPS4 dependent recognition specifies and restriction of bacterial growth in Arabidopsis. (A) Hypersensitive response (HR) assay in different *Arabidopsis* accessions using *Pseudomonas fluorescens* (*Pf*) Pf0-1 secreting AvrRps4 wild-type and structure-guided mutants. Constructs were delivered to Arabidopsis Ws-2 and *rrs1-1/rps4-21/rps4b-1* knock-out background and HR was recorded 20 hours post-infiltration. Fraction refers to number of leaves showing HR of 12 randomly inoculated leaves. This experiment was repeated at least three times with similar results. (B) In planta bacterial growth assays of *Pto* DC3000 secreting AvrRps4 wild-type and mutant constructs. Bacterial suspensions with OD_600_ = 0.001 were pressure infiltrated into the leaves of 4-5-week-old Arabidopsis plants. Values are plotted from three independent experiments (denoted in different colors). Statistical significance of the values was calculated by one-way ANOVA followed by post-hoc Tukey HSD analysis. Letters above the data points denotes significant differences (P<0.05). Detailed statistical summary can be found in Table 5.

**Table 5.** Tukey multiple comparisons of means at 95% family-wise confidence level Day 3 Col-0 Day 3 Ws-2

| **Construct** | **diff** | **lower bound** | **Upper bound** | **p adj** |
| --- | --- | --- | --- | --- |
| D164A-AvrRps4 | 0.80269085 | 0.36342384 | 1.24195786 | 0.0000053 |
| EE/AA-AvrRps4 | 1.07949193 | 0.64022492 | 1.51875894 | 0 |
| KRVY/AAAA-AvrRps4 | 1.14645988 | 0.70719287 | 1.58572689 | 0 |
| N171A-AvrRps4 | 0.49868711 | 0.0594201 | 0.93795413 | 0.0159246 |
| Q194A-AvrRps4 | 0.70408364 | 0.26481663 | 1.14335065 | 0.0001012 |
| EE/AA-D164A | 0.27680107 | -0.1624659 | 0.71606808 | 0.4599963 |
| KRVY/AAAA-D164A | 0.34376903 | -0.095498 | 0.78303604 | 0.2192971 |
| N171A-D164A | -0.3040037 | -0.7432708 | 0.13526327 | 0.3512228 |
| Q194A-D164A | -0.0986072 | -0.5378742 | 0.3406598 | 0.9873355 |
| KRVY/AAAA-EE/AA | 0.06696796 | -0.3722991 | 0.50623497 | 0.9979272 |
| N171A-EE/AA | -0.5808048 | -1.0200718 | -0.1415378 | 0.0025517 |
| Q194A-EE/AA | -0.3754083 | -0.8146753 | 0.06385872 | 0.1416124 |
| N171A-KRVY/AAAA | -0.6477728 | -1.0870398 | -0.2085058 | 0.0004724 |
| Q194A-KRVY/AAAA | -0.4423762 | -0.8816433 | -0.0031092 | 0.0472852 |
| Q194A-N171A | 0.20539652 | -0.2338705 | 0.64466353 | 0.759545 |

**Day 3 Ws-2**
| **Construct** | **diff** | **Lower bound** | **Upper bound** | **p adj** |
| --- | --- | --- | --- | --- |
| D164A-AvrRps4 | 1.26186508 | 0.7822084 | 1.74152177 | 0 |
| EE/AA-AvrRps4 | 1.2783785 | 0.7987218 | 1.75803519 | 0 |
| KRVY/AAAA-AvrRps4 | 1.12588572 | 0.646229 | 1.60554241 | 0 |
| N171A-AvrRps4 | 0.492187 | 0.0125303 | 0.97184369 | 0.0405978 |
| Q194A-AvrRps4 | 0.70761177 | 0.2279551 | 1.18726846 | 0.0004693 |
| EE/AA-D164A | 0.01651342 | -0.4631433 | 0.49617011 | 0.9999987 |
| KRVY/AAAA-D164A | -0.1359794 | -0.6156361 | 0.34367733 | 0.9644636 |
| N171A-D164A | -0.7696781 | -1.2493348 | -0.2900214 | 0.000099 |
| Q194A-D164A | -0.5542533 | -1.03391 | -0.0745966 | 0.0132372 |
| KRVY/AAAA-EE/AA | -0.1524928 | -0.6321495 | 0.32716391 | 0.9425012 |
| N171A-EE/AA | -0.7861915 | -1.2658482 | -0.3065348 | 0.0000643 |
| Q194A-EE/AA | -0.5707667 | -1.0504234 | -0.09111 | 0.0095929 |
| N171A-KRVY/AAAA | -0.6336987 | -1.1133554 | -0.154042 | 0.0025804 |
| Q194A-KRVY/AAAA | -0.418274 | -0.8979306 | 0.06138275 | 0.1263936 |
| Q194A-N171A | 0.21542477 | -0.2642319 | 0.69508146 | 0.7893133 |

In addition to Ws-2, we also performed a parallel set of experiments in Arabidopsis ecotype Col-0 (which encodes the RRS1-S allele) and the Col-0 *rrs1-3/rrs1b-1* (RRS1-S/RRS1B double knockout) line. Overall, we observed a weaker HR towards AvrRps4 wild-type and mutants in Col-0 in comparison to Ws-2. Nevertheless, a similar pattern of HR phenotypes were observed in Col-0 compared to Ws-2, and none of the AvrRps4 variants triggered HR in the Col-0 *rrs1-3/rrs1b-1* line (Fig. S5B). The pattern of HR phenotypes conferred by the AvrRps4 interface mutants further validates the AvrRps4^C^/RRS1^WRKY^ structure and the role of these residues in recognition of AvrRps4 by the RRS1/RPS4 receptor pair.

### Loss of HR correlates with bacterial growth in RRS1/RPS4-containing *A. thaliana*

Having demonstrated the role of AvrRps4 interface residues in effector-triggered HR in *A. thaliana*, we next investigated their effects on bacterial growth. We performed bacterial growth assays on Arabidopsis ecotypes Ws-2, Col-0, Ws-2 *rrs1-1/rps4-21/rps4b-1* and Col-0 *rrs1-3/rrs1b-1* (as detailed in the previous section) using *P. syringae* pv. *tomato (Pto)* DC3000 strain carrying AvrRps4 wild-type or each mutant. Since both the single mutants AvrRps4 E175A and E187A displayed the same impaired HR as the double AvrRps4 EE/AA mutant in our previous assays, we focused on AvrRps4 EE/AA mutant only for this assay. Bacterial growth was scored at three days post-infection (dpi). *Pto* DC3000 carrying wild-type AvrRps4 displayed reduced growth on Ws-2 when compared to the mutant background (Ws-2 *rrs1-1/rps4-21/rps4b-1*), presumably due to the activation of RRS1-R/RPS4-dependent immunity (Fig. 5B). The effector mutants AvrRps4 D164A, EE/AA, KRVY/AAAA, which displayed a complete loss of HR in Ws-2, show a severe or complete lack of restriction of bacterial growth in Ws-2 (Fig. 5B). *Pto* DC3000:AvrRps4 Q194A and *Pto* DC3000:AvrRps4 N171A showed reduced bacterial growth (but not full restriction) when compared to wild-type AvrRps4, even though they displayed a similar cell death phenotype in *N. tabacum* (albeit weaker for AvrRps4 Q194A) and HR in Arabidopsis (Fig. 4A, 5A). All the *Pto* DC3000:AvrRps4 variants tested displayed indistinguishable bacterial growth in RRS1-R/RPS4 loss of function line (Fig. 5B). Finally, all the *Pto* DC3000:AvrRps4 variants displayed similar bacterial growth profiles in Col-0 and Col-0 *rrs1-3/rrs1b-1* line when compared to Ws-2 and Ws-2 *rrs1-1/rps4-21/rps4b-1* (Fig. S5C).

### The RRS1B/RPS4B immune receptor pair displays similar recognition specificities towards AvrRps4 variants as RRS1/RPS4

In addition to RRS1/RPS4, the RRS1B/RPS4B pair can confer recognition of AvrRps4 in Arabidopsis (25). Sequence alignment reveals an overall 60% amino acid identity of the integrated WRKY domains from RRS1 and RRS1B, with the ‘WRKYGQK’ motif and all residues interfacing with AvrRps4^C^ conserved (Fig. S6). To explore AvrRps4 recognition by RRS1B/RPS4B, we performed ITC titrations of RRS1B^WRKY^ with wild-type AvrRps4^C^ in vitro. In comparison to RRS1^WRKY^, RRS1B^WRKY^ binds ∼3-fold more weakly to AvrRps4^C^ (Fig. S6). When comparing the binding kinetics to the strength of immune responses in planta, we observed a weaker RRS1B/RPS4B-dependent HR to AvrRps4 compared to RRS1/RPS4. Nonetheless, both NLR pairs displayed a similar profile of immune responses towards the AvrRps4 structure-guided mutants in transient cell death assays and in *A. thaliana* HR assays (Fig. S6).

## Discussion

Despite recent advances, structural knowledge of how diverse integrated domains in plant NLRs perceive pathogen effectors is limited. Here, we investigated how the integrated WRKY domain of the Arabidopsis NLR RRS1 binds to the *Pseudomonas* effector AvrRps4, and how this underpins RRS1/RPS4-dependent immunity in planta. Further, through this work, we gained insights into interfaces in the RRS1^WRKY^ domain that are crucial for perception of two structurally unrelated effectors from distinct bacterial pathogens, which may have implications for NLR integrated domain engineering.

Transcriptional reprogramming upon NLR activation is well established as an early immune response in plants (35–37), and direct interactions between NLRs and transcription factors have been reported (38–42). WRKY transcription factors are important molecular players in the regulation of plant growth and development, abiotic and biotic stresses (43–45). Typically, WRKY transcription factors target genes by binding W-box DNA in promoters, via a signature amino acid motif ‘WRKYGQK’, to either promote or repress transcription (46–49). As WRKY TFs play an important role in plant immunity, it is unsurprising that they are often found as integrated domains in NLR immune receptors (50), supporting the hypothesis that pathogen effectors enhance virulence by targeting WRKY transcription factors. Therefore, understanding how effectors bind to WRKY integrated domains may inform how effector/WRKY binding promotes disease. The structure of the AvrRps4^C^/RRS1^WRKY^ complex reveals that the effector directly interacts with the DNA binding ‘WRKYGQK’ motif, likely rendering it unavailable for binding to DNA. AvrRps4^C^ binds to *At*WRKY41 with similar thermodynamic parameters to RRS1^WRKY^, and interface mutants that prevent AvrRps4^C^ interaction with RRS1^WRKY^ prevent interaction with *At*WRKY41, supporting the hypothesis that AvrRps4^C^ binds different WRKYs via a similar interface. Therefore, we speculate that AvrRps4 binds WRKY transcription factors to sterically block their binding to DNA, promoting virulence. WRKY domain residues interacting with AvrRps4^C^ are well conserved in these transcription factors (Fig. S7), suggesting that AvrRps4 could target multiple WRKY domains to promote virulence. In addition to WRKY TFs, a recent publication suggests AvrRps4 can interact with BTS domains to affect pathogen colonization (51). Understanding whether these functions are related requires further investigation.

Comparing the AvrRps4^C^/RRS1^WRKY^ structure with that of the PopP2/RRS1^WRKY^ (30) reveals an overlapping binding site for the effectors, primarily mediated by the β2-β3 segment of WRKY domain. The second lysine of the ‘WRK^1^YGQK^2^’ motif (K^2^) is acetylated by PopP2, abolishing the affinity of WRKY domain for W-box DNA (13, 14, 30). Intriguingly, acetylation of K^2^ lysine by PopP2 abolished the association of AvrRps4 with RRS1^WRKY^ (13), highlighting the important role of this interface in mediating the association of RRS1^WRKY^ with both effectors. It also highlights the likely shared role of the effectors in preventing interaction of WRKY domains with DNA as their virulence activity, either via enzymatic modification or steric blocking.

Studies with the NLR pair Pik from rice have shown that the strength of effector binding to integrated domains in vitro can correlate with immune responses in planta (52–54). Of the AvrRps4 mutants we tested to validate the RRS1^WRKY^ interface, all except N171A and Q194A prevented binding in vitro (by ITC) and in planta (by co-IP), and these did not give cell death in *Nicotiana* species when co-expressed with either RRS1-R/RPS4 or RRS1-S/RPS4. Further, they did not give HR or restrict bacterial growth in Arabidopsis Ws-2 or Col-0 ecotypes (except for a partial restriction of bacterial growth for the D164A mutation in the Col-0 background). The N171A mutant retained the same level of binding as wild-type in vitro, and displayed the same in planta phenotypes, although restriction of bacterial growth in Arabidopsis was reduced compared to wild-type in both Ws-2 and Col-0 ecotypes. Finally, the Q194A mutant showed a reduced binding in vitro (∼7-fold compared to wild-type) but maintained an HR in Arabidopsis as well as displaying a restriction of bacterial growth in Arabidopsis, albeit reduced compared to wild-type. Interestingly, this mutant consistently showed a qualitative reduction in the intensity of cell death in *Nicotiana*. Taken together, these AvrRps4 mutations validate the complex with RRS1^WRKY^ in that they prevent interaction in vitro and in planta, but they are not sufficient to determine whether strength of binding in vitro can directly correlate with in planta phenotypes. Further studies, including additional mutants, will be required to study this in the RRS1/RPS4 system.

Structural studies of singleton NLRs have shown that interactions between effectors and multiple domains within an NLR can be essential for activation (55–58). It is yet to be established whether this is also the case for effector perception involving paired NLRs with integrated domains, although the rice blast pathogen effector AVR-Pia immunoprecipates with its sensor NLR Pia-2 (RGA5) when the integrated HMA domain has been deleted. However, this interaction does not promote immune responses in planta (59). Although unresolved in the structure of AvrRps4^C^ alone, or in complex with RRS1^WRKY^, the N-terminal KRVY motif is known to be required for both the virulence activity of the effector and its perception by RRS1/RPS4 (26, 27). Here, we verified that the quadruple mutant AvrRps4 KRVY/AAAA retains interaction with RRS1^WRKY^ at wild-type levels in vitro and in vivo, but did not trigger RRS1/RPS4-dependent responses in our in planta assays. This suggests that while binding of AvrRps4 to the RRS1^WRKY^ domain is essential for immune activation, an additional interaction mediated by the N-terminal region of the effector to a region of RRS1 and/or RPS4 outside this domain is also required for initiation of defence. Further studies are required to determine how additional receptor domains outside of integrated domains in NLR-IDs contribute to receptor function.

The Arabidopsis NLR pair RRS1B/RPS4B perceives AvrRps4, but not PopP2 (25). Phylogenetically, the RRS1 WRKY belongs to Group III of the WRKY superfamily, whereas RRS1B WRKY is grouped into Group IIe (14, 25, 49). Here we found that AvrRps4^C^ binds the RRS1B^WRKY^ with three-fold lower affinity and RRS1B/RPS4B shows a similar pattern of recognition specificity in planta but with reduced phenotypes compared to RRS1/RPS4. A full investigation addressing why AvrRps4 shows differential interaction strength and phenotypes between RRS1 and RRS1B is beyond the scope of this work, but will be a direction for future research.

The unique ability of RRS1/RPS4 to perceive two effectors that differ both in sequence and structure, via the same integrated domain, highlights the potential for engineering of sensor NLRs to recognize diverse effectors. Recently, the range of rice blast pathogen effectors recognized by the integrated HMA domain of Pia-2 (RGA5) has been expanded by molecular engineering (59). However, this expanded recognition was towards structurally related effectors and may not be via a shared interface. Further, although cell death responses were observed in *N. benthamiana*, the engineered NLR was not able to deliver an expanded disease resistance profile in transgenic rice. This suggests we still require a better understanding of how NLR-IDs interact with effectors, and their partner helper NLRs, to enable bespoke engineering of disease resistance.

## Materials and Methods

### Gene cloning

For in vitro studies, the gene fragments of AvrRps4^C^ (Gly134–Gln221), RRS1^WRKY^ (Ser1194-Thr1273), RRS1B^WRKY^ (Asn1164-Thr1241), *At*WRKY41 (Thr125-Ile204) were cloned in various pOPIN expression vectors using in-fusion cloning strategy as described in the *SI Materials and Methods*.

For transient assays in *N. tabacum* and *N. benthamiana*, domesticated genomic fragments encoding RRS1-R, RRS1-S, RRS1B, RPS4, RPS4B were cloned into binary vector pICSL86977 under a 35S (CaMV) promoter with C-terminal 6xHis/3xFLAG-tag using Golden Gate assembly method as described in (24). Similar cloning techniques were used to generate constructs expressing RRS1^WRKY+83^. Full length AvrRps4 (*P. syringae* pv. pisi) was PCR-amplified from published constructs (13, 24, 27) and assembled with a C-terminal 4xmyc-tag in binary vector pICSL86977 under the control of 35S (CaMV) promoter using Golden Gate assembly method. DNA encoding each mutation was synthesised and cloned into pICSL86977 as described above.

For HR and bacterial growth assays in *A. thaliana,* full-length AvrRps4 and variants were cloned into a golden-gate compatible pEDV3 vector with C-terminal 4xmyc-tag.

### Protein Production and Purification

Plasmids expressing in planta processed C-terminal fragment of AvrRps4 (AvrRps4^C^) and integrated WRKY domain of RRS1 (RRS1^WRKY^) was expressed in *E. coli* SHuffle cells. The proteins were purified via immobilized metal-affinity chromatography (IMAC) followed by size-exclusion chromatography. Purified fractions were pooled and concentrated to 15 mg/mL and used for further studies. Detailed procedures are provided in the *SI Materials and Methods*.

### Crystallization and Structure Determination

Crystals of the AvrRps4^C^/RRS1^WRKY^ complex were obtained from a 1:1 solution of 15 mg/mL protein with 0.8 M Potassium sodium tartrate tertrahydrate, 0.1 M Sodium HEPES pH 7.5. Diffraction data were collected at the Diamond Light Source on the i03 beamline and processed in P6_1/5_22 space group. The structure was determined by molecular replacement using the model of a monomer of AvrRps4^C^ (PDB ID: 4B6X) and the RRS1^WRKY^ from the PopP2/RRS1^WRKY^ complex (PDB ID: 5W3X) as search model. Further details are provided in the *SI Materials and Methods*. X-ray data collection and refinement statistics are summarized in Table 2.

### In vitro Protein–Protein Interaction studies

AvrRps4^C^/RRS1^WRKY^ complex formation in vitro was studied using analytical gel filtration chromatography and isothermal titration calorimetry (ITC). The effect of structure-guided mutations on the AvrRps4^C^/RRS1^WRKY^ interaction in vitro was investigated using isothermal titration calorimetry (ITC) as described in the *SI Materials and Methods*.

### Transient cell death assays and co-Immunoprecipitation studies

Agrobacterium mediated transient cell death assays were performed in *N. tabacum* and co-immunoprecpitation assays were performed in *N. benthamiana*. Detailed information concerning plant materials, growth conditions, plasmid construction and immunoblotting are provided in the *SI Materials and Methods*.

### Arabidopsis HR assays and bacterial growth assays

Bacterial strain *P. fluorescens* Pf0-EtHAn and *Pto*DC3000 was used for HR or in planta bacterial growth assays, respectively. The *Arabidopsis thaliana* accessions Ws-2 and Col-0 were used as wild-type for all the assays in this study. Further details about plant materials, growth conditions, plasmid construction and mobilization, pathogen infection assays and bacterial growth assays are provided in the *SI Materials and Methods*.

## Acknowledgments

This work was supported by the European Research Council [ERC; proposal 669926]; the UKRI Biotechnology and Biological Sciences Research Council (BBSRC) Norwich Research Park Biosciences Doctoral Training Partnership, UK [grant BB/M011216/1]; the UKRI BBSRC, UK [grants BB/P012574, BBS/E/J/000PR9795]; the BBSRC Future Leader Fellowship (grant BB/R012172/1). We would like to thank Dr. Clare Stevenson, Julia Mundy and Professor David Lawson from the JIC Biophysical Analysis and X-ray Crystallography platform for their support with ITC, CD spectroscopy, protein crystallization and X-ray data collection, Andrew Davies and Phil Robinson from JIC Scientific Photography for their help with leaf imaging. We also thank Dr. Kee Hoon Sohn for helpful suggestions for tri-parental mating and other members of Banfield and Jones laboratory for discussions.

## Supplementary information

### Materials and Methods

#### Protein production and purification

##### Gene cloning, expression, and purification of proteins for in vitro binding studies

Gene fragment of AvrRps4^C^ (134–221) was cloned in pOPIN-F (with a cleavable 6xHis-tag) expression vector while DNA fragments of RRS1^WRKY^ (1194–1273), RRS1B^WRKY^ (Asn1164-Thr1241), and *At*WRKY41 (Thr125-Ile204) were cloned in pOPIN-M (with a cleavable 6xHis-MBP-tag) expression vector using in-fusion cloning strategy (Clontech, Mountain View, CA, United States) (1). The constructs were then transformed in *Escherichia coli* (*E. coli*) SHuffle cells for expression. Bacterial cultures were grown in LB media (with 100 μg/mL carbenicillin) at 30°C to an OD_600_ = 0.6 followed by induction with 1 mM IPTG (isopropyl β-D-1-thiogalactopyranoside) and overnight growth 18°C. The cells were harvested by centrifugation at 6,000 g for 10 min and resuspended in Buffer A1 (50mM HEPES pH (8.0), 50mM glycine, 500mM NaCl, 30 mM imidazole and 5% v/v glycerol, EDTA free protease inhibitor tablets [1 tablet/50mL of A1 buffer]) followed by lysis by sonication with VC 750 VibraCell™ (Sonics) at 40 % amplitude, 1 sec on/3 sec off pulse for 20 min on ice. Cell debris was removed by centrifugation at 45,000 g for 60 mins. Purification of the proteins was performed using an ÄKTA Xpress purification system following two-step programme comprising initial capture by immobilised metal affinity chromatography (IMAC) [Step elution by Buffer B1 – Buffer A1 supplemented with 500 mM imidazole] followed by gel filtration with Superdex 75 26/600 gel filtration column pre-equilibrated in Buffer A4 (20 mM HEPES pH 7.5, 150 mM NaCl) supplemented with 1 mM TCEP. Fractions under the elution peak from the gel filtration columns were assessed by SDS-PAGE for the presence of the purified proteins and were then pooled and treated with 3C protease (10 µg/mg fusion protein) overnight at 4°C to cleave the 6xHis/6xHis-MBP-tag respectively. Cleaved 6xHis and 6xHis-MBP tags were then separated from their respective digested protein samples by passing the samples through a Ni^2+^-NTA column and collecting the flow-through and wash samples. These samples were assessed by SDS-PAGE, pooled and concentrated via 3 kDa cut-off spin concentrators before subjecting to a second round of size-exclusion with a Superdex 75 16/600 gel filtration column pre-equilibrated in Buffer A4. Eluted samples were then concentrated via 3 kDa cut-off spin concentrators to a final concentration of 10-15 mg/mL (as calculated by Nanodrop (Thermofisher Scientific™, NanoDrop™ One Microvolume UV-Vis Spectrophotometer) at A_205_) and were aliquoted and flash frozen at −80°C for subsequent analysis.

##### Expression and purification of proteins for crystallization

For crystallization of the AvrRps4^C^/RRS1^WRKY^ complex, AvrRps4^C^ was cloned in pOPIN-A to express untagged protein and pOPIN-M construct of RRS1^WRKY^ (with cleavable 6xHis-MBP-tag) was used as mentioned above. Both the constructs were transformed individually into *E. coli* SHuffle cells and expressed using the above-mentioned conditions. The bacterial cells expressing AvrRps4^C^ and RRS1^WRKY^ were then mixed together, lysed and the proteins were co-purified via IMAC followed by size-exclusion chromatography as mentioned above. Eluted fractions were assessed by SDS-PAGE. The presence of untagged AvrRps4^C^ in the RRS1^WRKY^ eluted fractions confirmed complex formation in vitro. The eluted complex was subjected to 3C protease cleavage overnight at 4°C followed by an IMAC step to remove the cleaved 6xHis-MBP-tag. Fractions containing the untagged complex were then pooled, concentrated, and subjected to final gel filtration chromatography. The purified complex was concentrated to 15 mg/mL, aliquoted and used for crystallization studies.

##### Crystallization, data collection and structure solution

For crystallization of the AvrRps4^C^/RRS1^WRKY^ complex, the sitting drop vapour diffusion method was used. Potential crystallization conditions were explored using commercially available crystallization screens. All crystallization trials were setup in 96-well plates, using an Oryx nano robot (Douglas Instruments) at a concentration of 7.5 mg/mL and 15 mg/mL at 20°C. Crystals of the AvrRps4^C^/RRS1^WRKY^ complex appeared after few weeks in a condition comprising 0.8M Potassium sodium tartrate tertrahydrate, 0.1 M Sodium HEPES pH 7.5 from the Morpheus^TM^ screen. The crystals were snap frozen in liquid nitrogen and shipped to the Diamond Light Source for X-ray data collection.

Diffraction data was collected at Diamond Light Source, i03 beamline, under proposal mx18565. The data were scaled and merged by Aimless in the CCP4i2 software package (2). The AvrRps4^C^/RRS1^WRKY^ complex structure was solved by molecular replacement using PHASER (3) with the structures of AvrRps4^C^ (PDB ID: 4B6X) and PopP2/RRS1^WRKY^ (PDB ID: 5W3X) as search models. Iterative cycles of manual model building using COOT (4) and ISOLDE (5) and refined using REFMAC (6) produced the final structure, which was then validated using MolProbity (7). Interaction interfaces were analyzed using PdbePISA (8). Models were visualized using ChimeraX (9). The final protein model, and the data used to derive it, can be found in Protein Data Bank (PDB) (https://www.ebi.ac.uk/pdbe/) with the PDB ID: 7P8K.

##### Circular dichroism spectroscopy

Purified AvrRps4^C^ wild-type and mutants were dialyzed in 10mM phosphate buffer, pH 8.0 at a final concentration of 0.5 mg/mL. Samples were analyzed in the far-UV region between 190-260 nm at 20°C by using Chirascan™ plus CD Spectrometer (Applied Photophysics) and quartz cuvette of path length 1mm. For each sample, three successive spectral scans were averaged and adjusted by subtracting corresponding blanks. The results were plotted using ggplot2 in R (10)

##### In vitro Protein–Protein Interaction studies Analytical gel filtration

To study AvrRps4^C^ and RRS1^WRKY^ complex formation in vitro, individual proteins (at a concentration of 1 mg/mL) were applied to pre-equilibrated (Equilibration buffer - 20 mM HEPES pH 7.5, 150 mM NaCl, 1 mM TCEP) Superdex 75 10/300 analytical column (GE-Healthcare) using an AKTA Explorer (GE-Healthcare) at 4°C and eluted at a flow rate of 0.5 mL/min by monitoring the absorbance at 280 nm. 500 μL fractions were collected and analyzed by SDS-PAGE. For complex formation, proteins were combined in a 1:1 molar ratio and incubated on ice for 1-2 hrs before the analysis. The results were plotted using ggplot2 in R (10)

##### Isothermal titration calorimetry (ITC)

ITC experiments were performed using a MicroCal PEAQ-ITC (Malvern, UK). To test the interaction of AvrRPS4^C^ wild-type or structure-guided mutants with RRS1^WRKY^, *At*WRKY41 or RRS1B^WRKY^, the calorimetric cell was filled with 20 μM of RRS1^WRKY^/*At*WRKY 41/RRS1B^WRKY^ and titrated with 200 μM of AvrRps4^C^ wild-type/mutants in the syringe. Each ITC run included a single injection of 0.5 μL followed by 18 injections of 2 μL each. Injections were made at 120-second intervals with a stirring speed of 750 rpm. Data were processed with AFFINImeter ITC analysis software (11). ITC runs for wild-type and mutants were done in triplicate at 25°C using buffer A4. All the ITC curves were plotted using ggplot2 in R (10).

### Transient cell death assays and co-Immunoprecipitation studies

#### *N. tabacum* cell death assays

Transient cell death assays were performed using 4-5 week-old *N. tabacum* “Petit Gerard” as described previously (12). Plants were grown in long days (16 hr light/8 hr dark) under high light intensity at 24°C. *Agrobacterium tumefaciens* GV3101 was used to deliver C-terminal 4xmyc-tagged full-length constructs of AvrRps4 wild-type and mutants, and C-terminal 6xHis/3xFLAG-tagged RRS1-R, RRS1-S, RRS1B, RPS4, RPS4B. Agrobacterium cells expressing these constructs were grown at 28°C, harvested, and resuspended in infiltration buffer (10 mM MgCl_2_, 10 mM MES [pH 5.6]), supplemented with 150 μM acetosyringone. Appropriate combinations of the above constructs were mixed at an OD_600_ = 0.5 per construct and were hand infiltrated on the abaxial surface of 4-5 week-old *N. tabacum* leaves by a 1ml needleless syringe. Infiltrated leaves were detached 5 days post infiltration (dpi) and imaged under white light. The experiment was done in triplicate with similar results.

#### In planta co-immunoprecipitation assays

For co-immunoprecipitation assays, proteins were transiently expressed in 4-5 week old *N. benthamiana* leaves using agroinfiltration as described in (13). Leaf samples (4 g) were harvested at 3 dpi, frozen in liquid nitrogen, and ground to fine powder. A total of 8 mL (two times weight/volume) of ice-cold protein extraction buffer [10% glycerol, 1 mM EDTA, 25 mM Tris [pH 7.5], 150 mM NaCl, 2% w/v PVPP, 10 mM DTT, 1x protease inhibitor cocktail [Sigma], 1 % vol/vol Nonidet P-40] was added to the ground tissue and samples were centrifuged at 6,000 × g at 4°C for 15 min. Supernatant was then filtered through miracloth to remove residual plant debris. 50μL of the filtrate was aliquoted and ran on 4-20 % precast SDS-PAGE gels to check for the expression of the proteins in the input fraction. Residual samples were mixed with 50 μL Flag beads and incubated at 4°C (with constant rotation) for 1 hr. Flag beads were washed three times with IP buffer (10% glycerol, 1 mM EDTA, 25 mM Tris [pH 7.5], 150 mM NaCl, 1 % vol/vol Nonidet P-40) and re-suspended in 30 μL SDS-loading buffer. Immunoprecipitated samples were recovered from the flag beads by boiling at 70°C for 10 min. Eluted samples were separated by 4-20% precast SDS-PAGE, electroblotted onto PVDF membranes (Bio-Rad), and probed with HRP-conjugated anti-FLAG M2 (1:10000 dilution, Sigma) and anti-Myc (1:5000, Santa Cruz) as required.

#### Arabidopsis HR assays and bacterial growth assays Plant material and growth conditions

Arabidopsis accessions Ws-2 and Col-0 were used as wild-type for all the assays in this study. Ws-2 was the background of the triple mutant *rrs1-1*/*rps4-21*/*rps4b-1* while double mutant *rrs1-3*/*rrs1b-1* and single mutants *rrs1-3* and *rrs1b-1* lines were in the Col-0 background. The plants were grown under short day conditions (10-h light/14-h dark) at 22°C and 65% humidity for 4-5 weeks before being used for assays.

#### Arabidopsis HR assays

For Arabidopsis HR assays, Pf0-EtHAn was grown on King’s B agar medium containing chloramphenicol (30 μg/ mL) at 28°C. Plasmids were mobilized into Pf0-EtHAn using tri-parental mating method using *E. coli* HB101 (pRK2013) as a helper strain as described in (14). For HR assays bacteria were grown overnight at 28°C in KB media and cells were harvested, washed and resuspended in freshly prepared sterile 10 mM MgCl_2_. Final concentration of the inoculum was adjusted to OD_600_ = 0.3. The inoculum was then hand infiltrated into the leaves of 5-week-old plants with 1-ml needleless syringes. 5-6 leaves were infiltrated for each construct per genotype per biological replicate. Plants were then blotted with the tissue to remove the excess bacteria and then kept at 22°C covered with a transparent dome. HR was scored 20 hrs post infection. The experiment was repeated three times with similar results.

#### Bacterial growth assay

Pto DC3000 containing full-length 4xc-myc-tagged wild-type AvrRps4, or structure guided AvrRps4 mutant variants and the AvrRps4 KRVY/AAAA mutant (negative control) were grown on selective King’s B (KB) medium plates (containing 50 μg/mL Rifampicin and 20 μg/mL Gentamycin) for 48 h at 28°C. Bacterial cells were harvested, washed and resuspended in sterile 10 mM MgCl_2_ to a final OD_600_ = 0.001. The bacterial suspension was then hand infiltrated on the abaxial surface of 5-week-old Arabidopsis leaves using a 1mL needleless syringe. For the bacterial growth assays, 2 leaves each of 10 independent plants/genotype/construct constitute one biological replicate with three replicates in total. Samples from 4 plants were collected at day 0 and samples from 6 plants were collected at day 3. For quantification, 2 leaf discs from one plant (one leaf disc per leaf) were collected with a 6-mm-diameter cork borer (disc area −0.283 cm^2^) and were ground in 200 μL of infiltration buffer (10 mM MgCl_2_). For day 0, samples from 4 plants were independently ground and spotted (10 μL /spot) on selective KB medium. For day 3, samples from 6 plants were independently ground, serially diluted (5, 50, 5X10^2^, 5X10^3^ and 5X10^4^ times) and spotted (6 μL/spot) on selective KB medium. The plates were incubated at 28°C for two days before colony forming units (CFU/drop) were calculated. Bacterial growth is represented as CFU cm^-2^ of leaf tissue. Statistical significance was determined by one-way ANOVA followed by post-hoc Tukey HSD analysis. The results were plotted using ggplot2 in R (10).

---

**Fig. S1.**
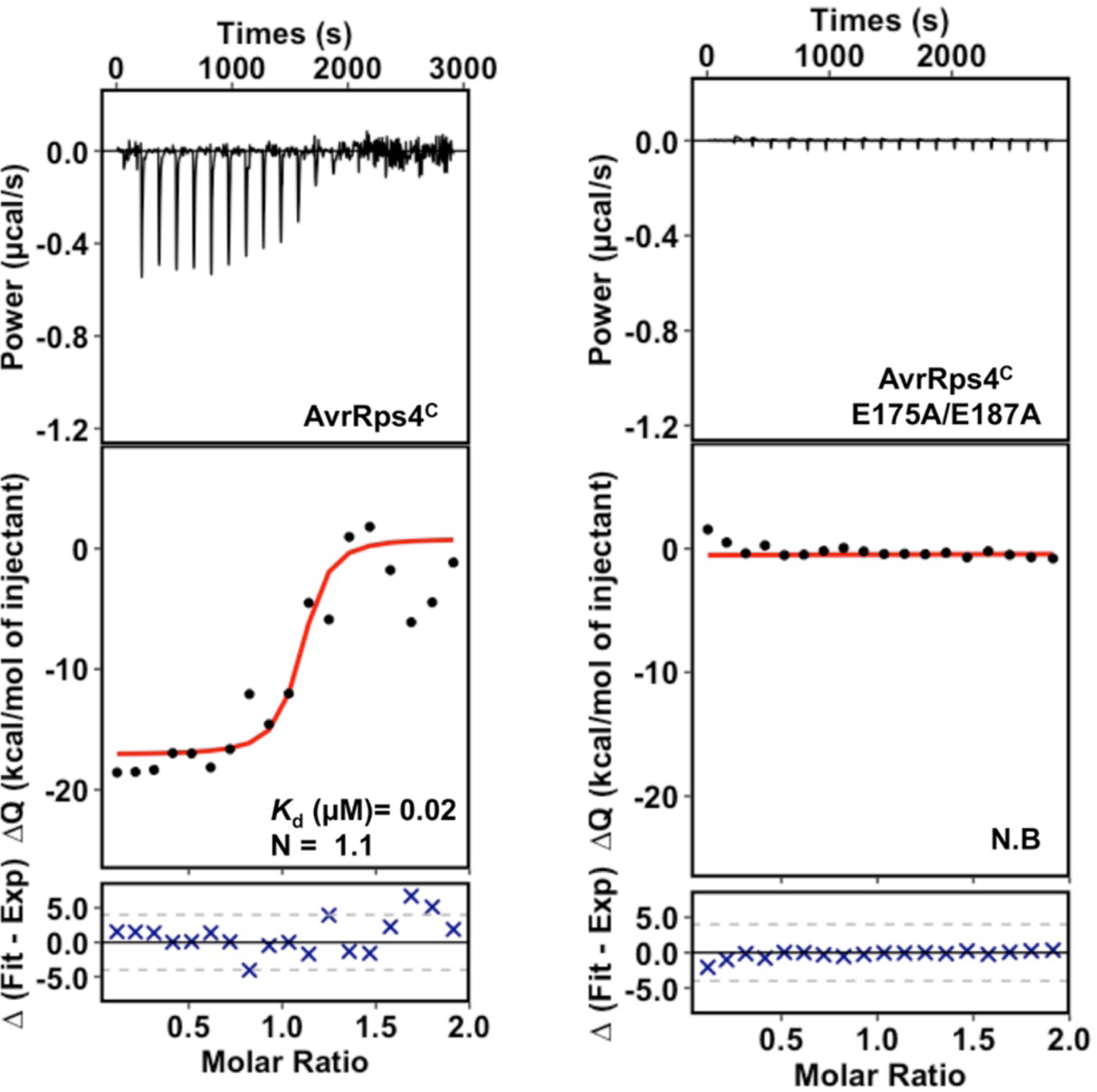
AvrRps4^C^ interacts with *At*WRKY41 in vitro. Isothermal titration calorimetry (ITC) of *At*WRKY41 with wild-type AvrRps4^C^ and AvrRps4 E175A/E187A (EE/AA) mutant. Raw processed thermogram after baseline correction and noise removal is displayed in the upper panel. The lower panel represents the experimental binding isotherm obtained for the interaction of AvrRps4^C^ and mutant with *At*WRKY41 together with the global fitted curves (displayed in red) obtained from three independent experiments using AFFINImeter software (11). The *K*_d_ was derived from fitting to a 1:1 binding model.

**Fig. S2.**
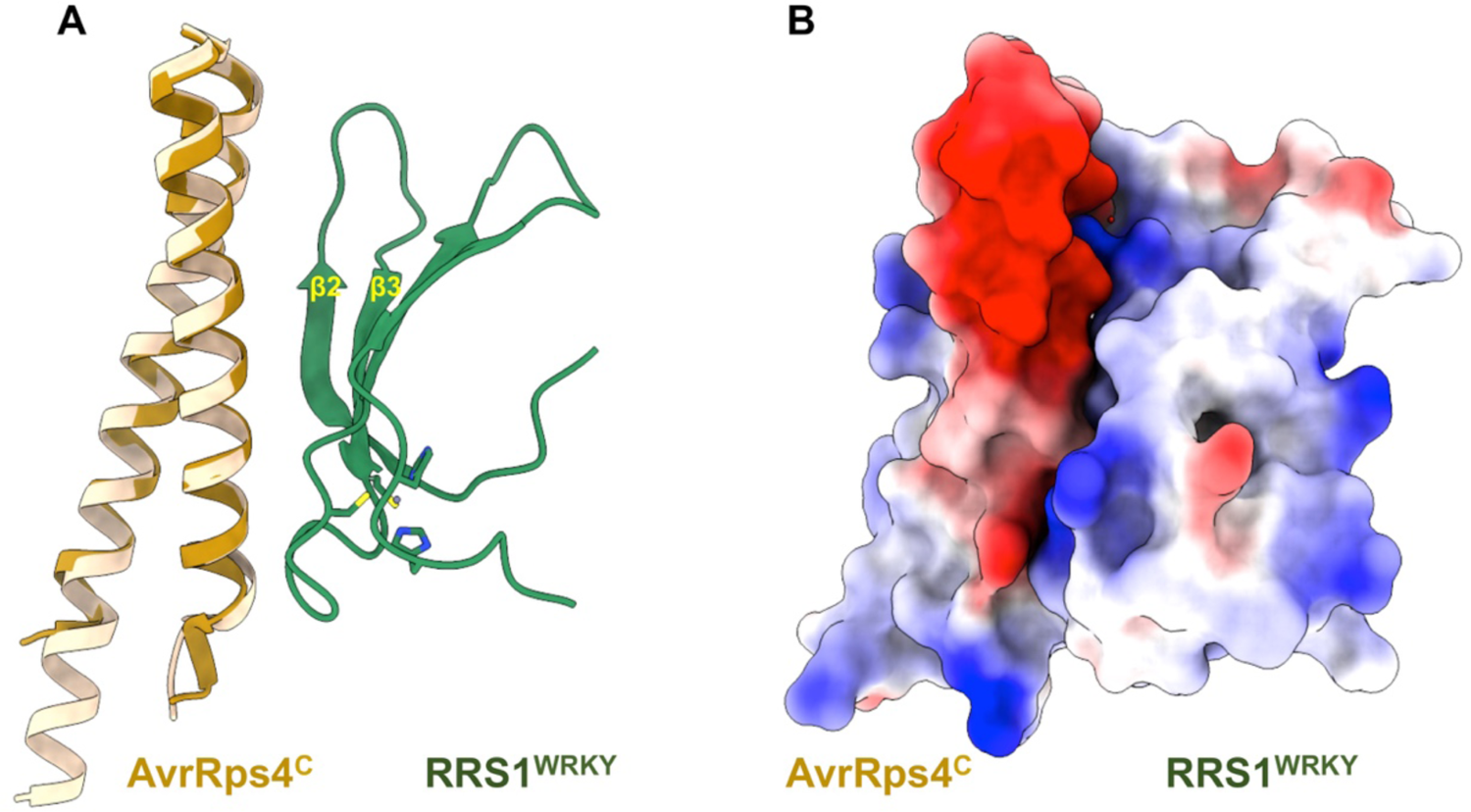
(A) Overlay of the crystal structure of AvrRps4^C^/RRS1^WRKY^ (Dark goldenrod/Dark green) with the previously published crystal structure of AvrRps4^C^ (light goldenrod) (PDB ID: 4B6X). (B) Electrostatic surface representation of the AvrRps4^C^/RRS1^WRKY^ complex highlighting the electronegative patch in AvrRps4^C^ and electropositive patch in RRS1^WRKY^ at the interface.

**Fig. S3.**
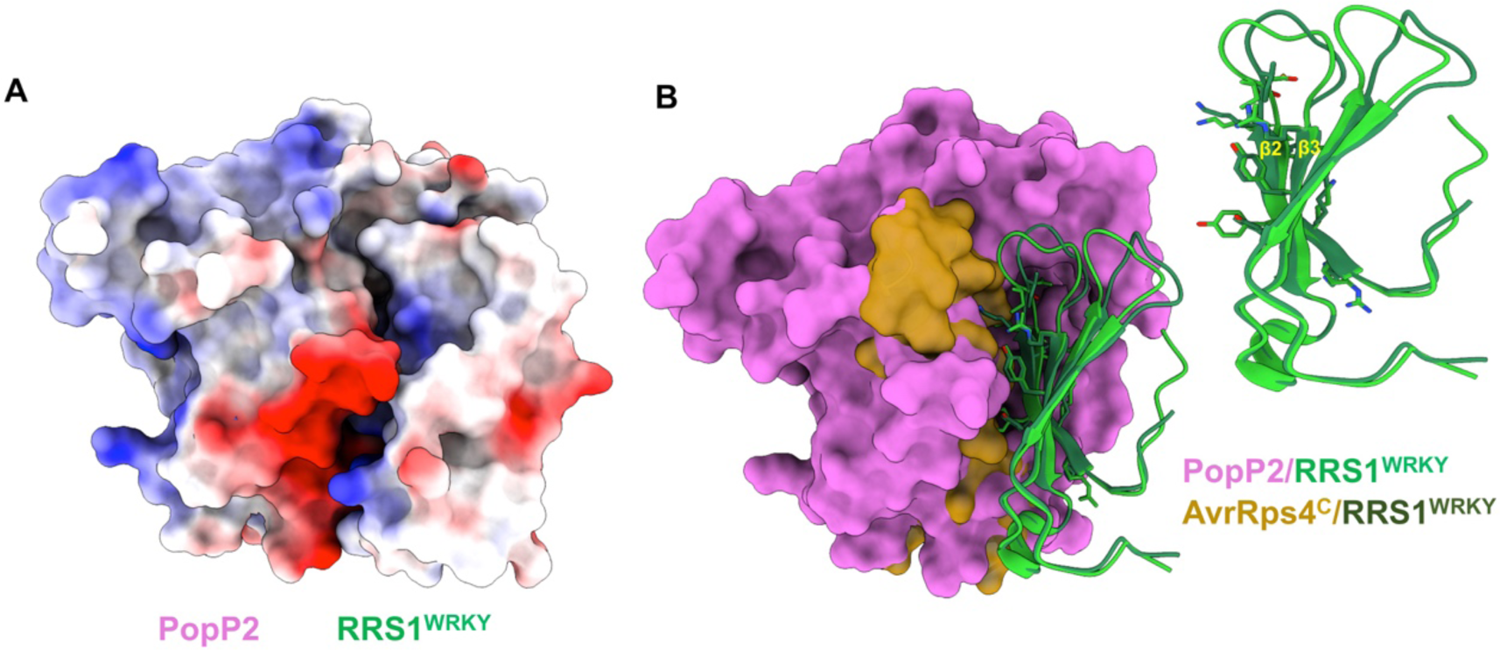
(A) Electrostatic surface representation of the PopP2/RRS1^WRKY^ complex structure (PDB ID: 5W3X) highlighting the electronegative patch on PopP2 and electropositive patch in RRS1^WRKY^ at the interface. (B) Overlay of the crystal structure of AvrRps4^C^/RRS1^WRKY^ (gold (surface)/green (ribbon) with PopP2/RRS1^WRKY^ (PDB ID: 5W3X, purple (surface)/green ribbon) based on the WRKY domains (left). Comparative view of β2, β3 segments of RRS1^WRKY^ mediating the interaction with AvrRps4^C^ (Dark green) and PopP2 (light green) is displayed (right).

**Fig. S4.**
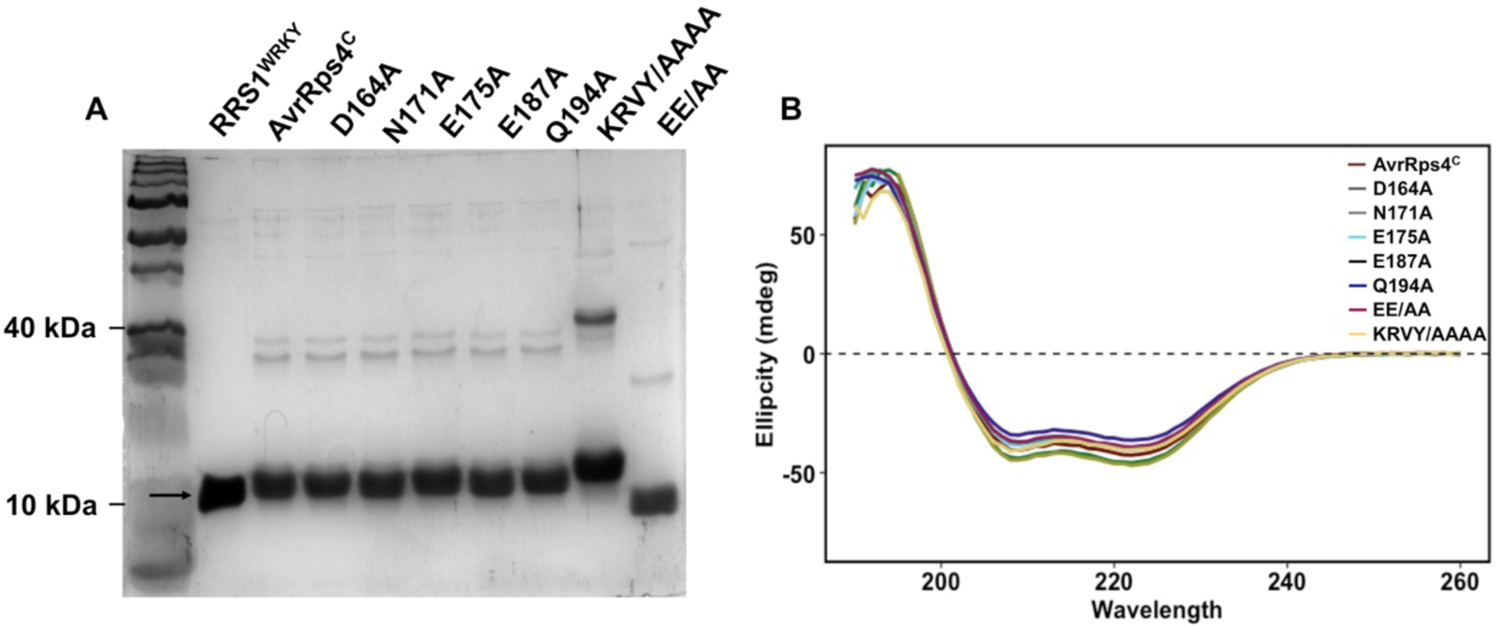
(A) SDS-PAGE of RRS1^WRKY^ and AvrRps4^C^ (wild-type and mutants) samples used for in vitro studies. Arrow indicates the expected size of the purified proteins. (B) CD spectra of the wild-type AvrRps4^C^ and mutants. Far-UV spectra corresponding to the wild-type (brick red), D164A (dark green), N171A (olive green), E175A (cyan), E187A (bluish green), Q194A (dark blue), EE/AA (purple) and KRVY/AAAA (coral) are shown. Spectra were taken at 20°C using 0.5 mg/mL of each protein. Each scan represents the average of three independent measurements.

**Fig. S5.**
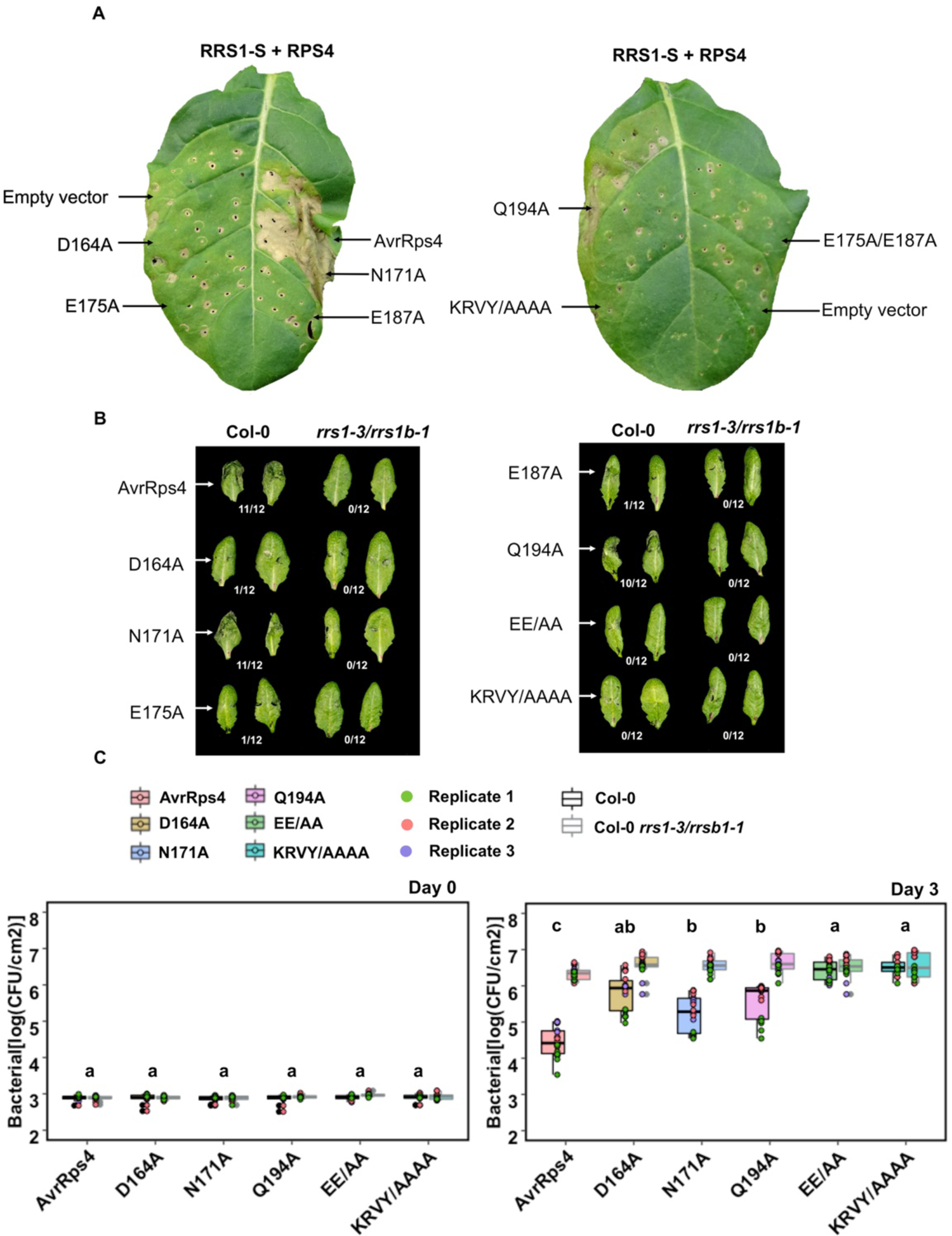
Structural guided mutants of AvrRps4 confer distinct RRS1-S/RPS4 dependent recognition specifies. (A) Structure-guided mutants in AvrRps4 at the AvrRps4^C^/RRS1^WRKY^ interface compromise RRS1-S/RPS4 mediated cell death in *N. tabacum*. Representative leaf images show RRS1-S/RPS4 mediated cell death response to wild-type and a subset of structure-guided mutants of AvrRps4. Agroinfiltration assays were performed in 4-5-week-old *N. tabacum* leaves, and cell death was assessed at 4 dpi. The experiment was repeated three times with similar results. (B) Hypersensitive response (HR) assay in Arabidopsis lines using *Pseudomonas fluorescens* (*Pf*) Pf0-1 secreting AvrRps4 wild-type and mutants. Constructs were delivered from (*Pf*) Pf0-1 into Arabidopsis Col-0 and Col-0 *rrs1-3/rrsb-1* background and HR observed 20 hours post-infiltration. Fraction refers to number of leaves showing HR of 12 randomly inoculated leaves. This experiment was repeated at least three times with similar results. (C) In planta bacterial growth assays with Pto DC3000 secreting AvrRps4 wild-type and mutants on Col-0 and Col-0 *rrs1-3/rrsb-1* background plants. Bacterial suspensions with OD_600_ = 0.001 were pressure infiltrated into the leaves of 5-week-old Arabidopsis plants. Values are plotted from three independent experiments (shown in different colors). Statistical significance of the values was calculated by one-way ANOVA followed by post-hoc Tukey HSD analysis. Letters above the data points denotes significant differences (P<0.05). Detailed statistical summary can be found in Table 5.

**Fig. S6.**
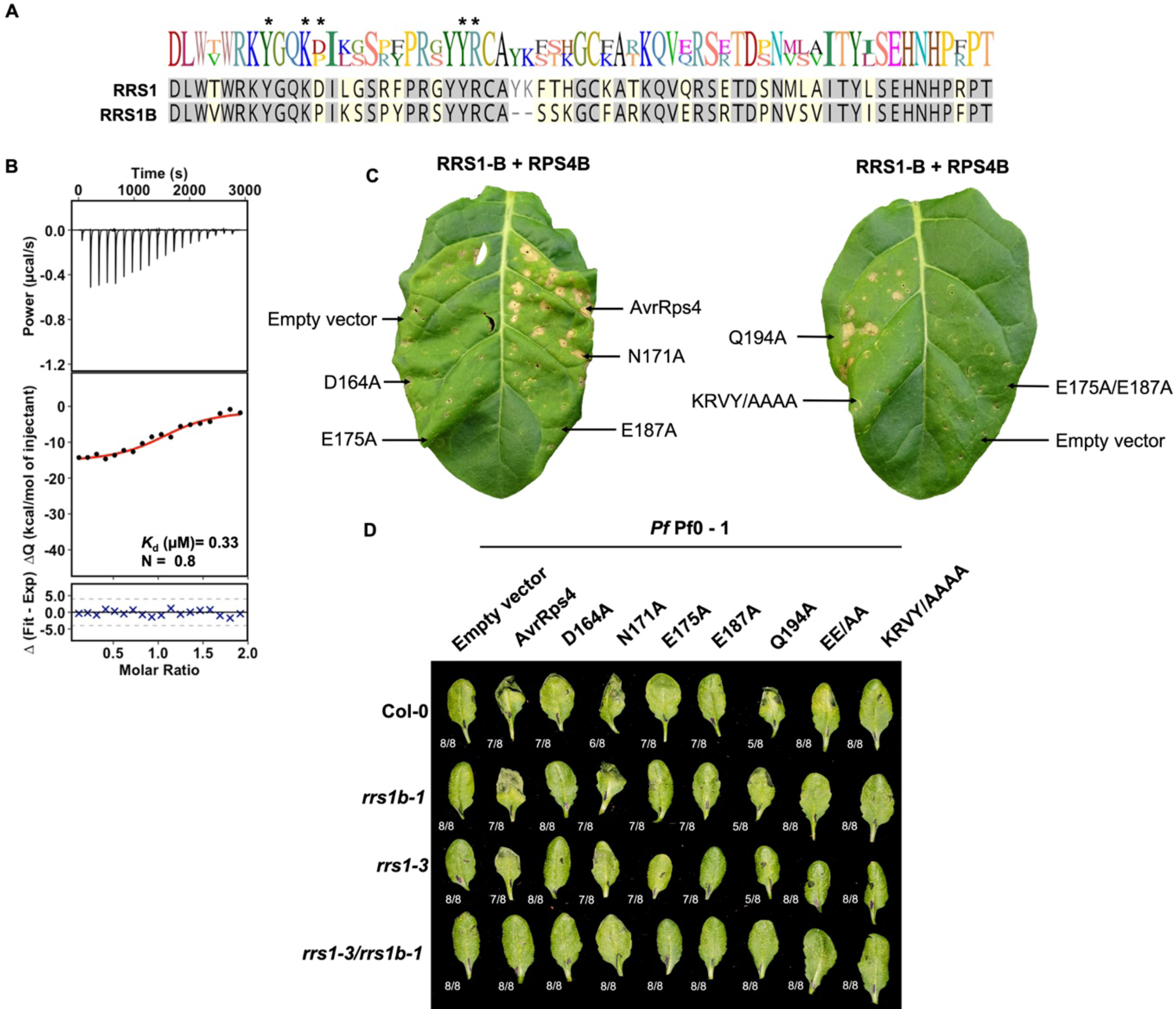
(A) Protein sequence alignment of the integrated WRKY domains from RRS1 and RRS1B. Grey shaded letters show identical residues. Residues at the AvrRps4^C^/RRS1^WRKY^ interface are marked with asterisk. (B) Isothermal titration calorimetry (ITC) titrations of RRS1B^WRKY^ with wild-type AvrRps4^C^. Raw processed thermogram after baseline correction and noise removal is displayed in the upper panel. The lower panel shows the experimental binding isotherm for the interaction together with the global fitted curve (displayed in red) obtained from three independent experiments using AFFINImeter software (11). The *K*_d_ was derived from fitting to a 1:1 binding model. (C) Structural guided mutants of AvrRps4 at the AvrRps4^C^/RRS1^WRKY^ interface compromise RRS1B/RPS4B mediated cell death responses in *N. tabacum*. Representative leaf images show RRS1B/RPS4B mediated cell death response to AvrRps4 wild-type and a subset of structure-guided mutants. Agroinfiltration assays were performed in 4- to 5-week-old *N. tabacum* leaves, and HR phenotypes were assessed at 4 dpi. The experiment was repeated three times with similar results. (D) Hypersensitive response (HR) assay in different *Arabidopsis* lines using *Pseudomonas fluorescens* (*Pf*) Pf0-1 secreting AvrRps4 wild-type and mutants. Constructs were delivered from Pf0-1 into Arabidopsis Col-0, Col-0 *rrs1-3*, Col-0 *rrs1b-1*, Col-0 *rrs1-3/rrs1b-1* background and HR was observed 20 hours post-infiltration. Fraction refers to number of leaves showing HR of 8 randomly inoculated leaves. This experiment was repeated twice with similar results.

**Fig. S7.**
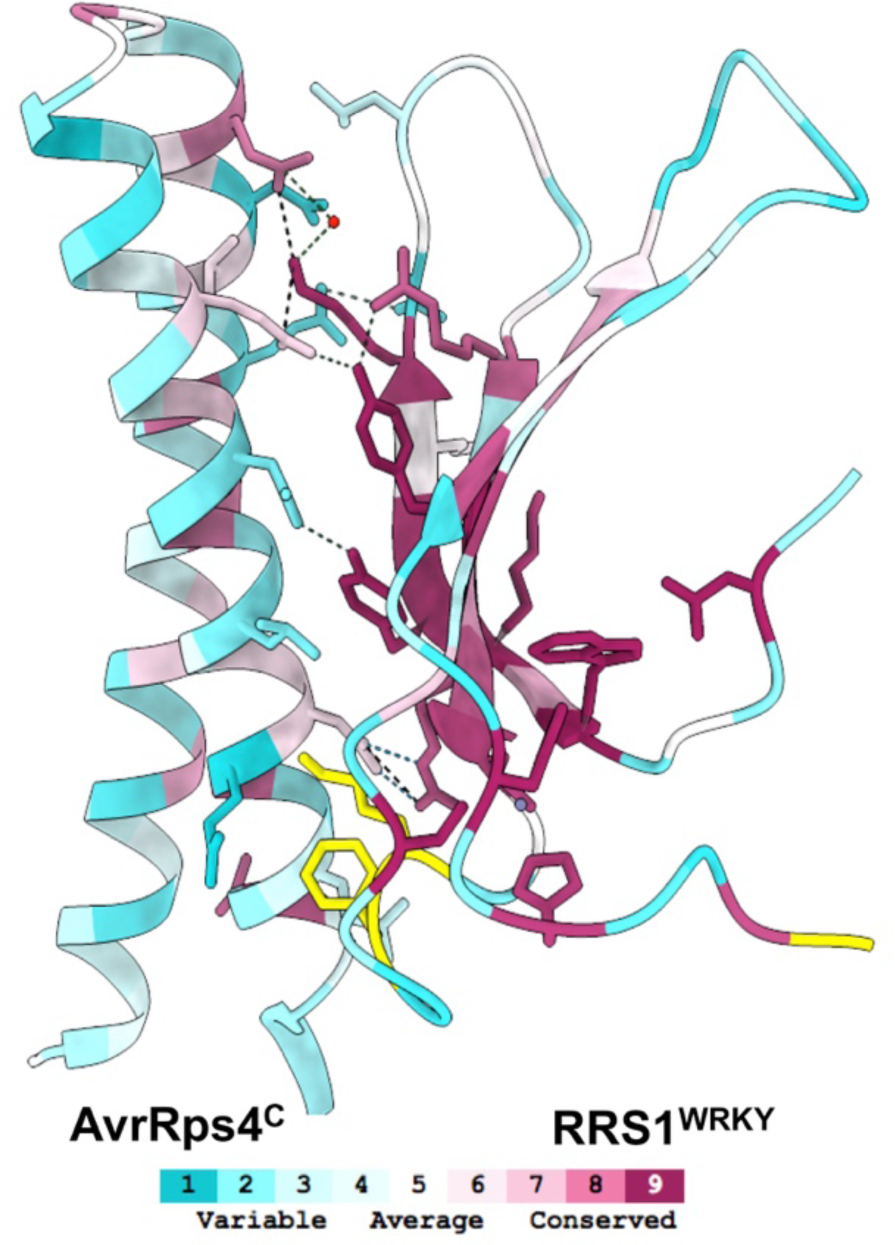
ConSurf analysis for the interface of the AvrRps4^C^/RRS1^WRKY^ complex. The conservation profiles of residues mediating the interaction of AvrRps4^C^ and RRS1^WRKY^ as calculated by Consurf are mapped upon the structures (15). The complex is shown in cartoon representation and residues mediating interaction at the interface are highlighted. Each segment of the cartoon has been colored according to its conservation status ranging from purple (highly conserved) through white (moderately conserved) to cyan (highly variable). Segments highlighted in yellow are residues for which a meaningful conservation level could not be derived from the set of homologues sequences used for the analysis.

## References

1. P. N. Dodds, J. P. Rathjen, Plant immunity: towards an integrated view of plant– pathogen interactions. Nature Reviews Genetics 11, 539–548 (2010).

2. N. Mukhi, D. Gorenkin, M. J. Banfield, Exploring folds, evolution and host interactions: understanding effector structure/function in disease and immunity. New Phytologist 227, 326–333 (2020).

3. M. Khan, D. Seto, R. Subramaniam, D. Desveaux, Oh, the places they’ll go! A survey of phytopathogen effectors and their host targets. The Plant Journal 93, 651–663 (2018).

4. J. D. G. Jones, J. L. Dangl, The plant immune system. Nature 444, 323–329 (2006).

5. A. R. Bentham et al., A molecular roadmap to the plant immune system. Journal of Biological Chemistry 295, 14916–14935 (2020).

6. J. D. G. Jones, R. E. Vance, J. L. Dangl, Intracellular innate immune surveillance devices in plants and animals. Science 354, aaf6395 (2016).

7. H. H. Flor, Current Status of the Gene-For-Gene Concept. Annual Review of Phytopathology 9, 275–296 (1971).

8. A. C. Barragan, D. Weigel, Plant NLR diversity: the known unknowns of pan-NLRomes. The Plant Cell 33, 814–831 (2021).

9. Z. Duxbury et al., Pathogen perception by NLRs in plants and animals: Parallel worlds. Bioessays 38, 769–781 (2016).

10. S. Cesari, M. Bernoux, P. Moncuquet, T. Kroj, P. N. Dodds, A novel conserved mechanism for plant NLR protein pairs: the “integrated decoy” hypothesis. Frontiers in plant science 5, 606–606 (2014).

11. Marc T. Nishimura, F. Monteiro, Jeffery L. Dangl, Treasure Your Exceptions: Unusual Domains in Immune Receptors Reveal Host Virulence Targets. Cell 161, 957–960 (2015).

12. A. Maqbool et al., Structural basis of pathogen recognition by an integrated HMA domain in a plant NLR immune receptor. eLife 4, e08709 (2015).

13. Panagiotis F. Sarris et al., A Plant Immune Receptor Detects Pathogen Effectors that Target WRKY Transcription Factors. Cell 161, 1089–1100 (2015).

14. C. Le Roux et al., A Receptor Pair with an Integrated Decoy Converts Pathogen Disabling of Transcription Factors to Immunity. Cell 161, 1074–1088 (2015).

15. S. Cesari et al., The Rice Resistance Protein Pair RGA4/RGA5 Recognizes the Magnaporthe oryzae Effectors AVR-Pia and AVR1-CO39 by Direct Binding The Plant Cell 25, 1463-1481 (2013).

16. S. Cesari, M. Bernoux, P. Moncuquet, T. Kroj, P. N. Dodds, A novel conserved mechanism for plant NLR protein pairs: the “integrated decoy” hypothesis. Front Plant Sci 5, 606 (2014).

17. H. Adachi, L. Derevnina, S. Kamoun, NLR singletons, pairs, and networks: evolution, assembly, and regulation of the intracellular immunoreceptor circuitry of plants. Curr Opin Plant Biol 50, 121–131 (2019).

18. S. U. Huh et al., Protein-protein interactions in the RPS4/RRS1 immune receptor complex. PLOS Pathogens 13, e1006376 (2017).

19. M. Narusaka, K. Hatakeyama, K. Shirasu, Y. Narusaka, Arabidopsis dual resistance proteins, both RPS4 and RRS1, are required for resistance to bacterial wilt in transgenic Brassica crops. Plant Signal Behav 9, e29130 (2014).

20. W. Gassmann, M. E. Hinsch, B. J. Staskawicz, The Arabidopsis RPS4 bacterial-resistance gene is a member of the TIR-NBS-LRR family of disease-resistance genes. The Plant Journal 20, 265–277 (1999).

21. M. Hinsch, B. Staskawicz, Identification of a new Arabidopsis disease resistance locus, RPs4, and cloning of the corresponding avirulence gene, avrRps4, from Pseudomonas syringae pv. pisi. Mol Plant Microbe Interact 9, 55-61 (1996).

22. L. Deslandes et al., Physical interaction between RRS1-R, a protein conferring resistance to bacterial wilt, and PopP2, a type III effector targeted to the plant nucleus. Proceedings of the National Academy of Sciences 100, 8024 (2003).

23. C. Tasset et al., Autoacetylation of the Ralstonia solanacearum Effector PopP2 Targets a Lysine Residue Essential for RRS1-R-Mediated Immunity in Arabidopsis. PLOS Pathogens 6, e1001202 (2010).

24. Y. Ma et al., Distinct modes of derepression of an <em>Arabidopsis</em> immune receptor complex by two different bacterial effectors. Proceedings of the National Academy of Sciences 115, 10218 (2018).

25. S. B. Saucet et al., Two linked pairs of Arabidopsis TNL resistance genes independently confer recognition of bacterial effector AvrRps4. Nature Communications 6, 6338 (2015).

26. K. H. Sohn, Y. Zhang, J. D. Jones, The Pseudomonas syringae effector protein, AvrRPS4, requires in planta processing and the KRVY domain to function. Plant J 57, 1079-1091 (2009).

27. K. H. Sohn, R. K. Hughes, S. J. Piquerez, J. D. G. Jones, M. J. Banfield, Distinct regions of the <em>Pseudomonas syringae</em> coiled-coil effector AvrRps4 are required for activation of immunity. Proceedings of the National Academy of Sciences 109, 16371 (2012).

28. J. Su et al., The Conserved Arginine Required for AvrRps4 Processing Is Also Required for Recognition of Its N-Terminal Fragment in Lettuce. Molecular Plant-Microbe Interactions® 34, 270–278 (2020).

29. M. K. Halane et al., The bacterial type III-secreted protein AvrRps4 is a bipartite effector. PLOS Pathogens 14, e1006984 (2018).

30. Z.-M. Zhang et al., Mechanism of host substrate acetylation by a YopJ family effector. Nat Plants 3, 17115–17115 (2017).

31. M. S. Mukhtar et al., Independently Evolved Virulence Effectors Converge onto Hubs in a Plant Immune System Network. Science 333, 596 (2011).

32. M.-R. Duan et al., DNA binding mechanism revealed by high resolution crystal structure of Arabidopsis thaliana WRKY1 protein. Nucleic Acids Res 35, 1145–1154 (2007).

33. E. Krissinel, K. Henrick, Inference of Macromolecular Assemblies from Crystalline State. Journal of Molecular Biology 372, 774–797 (2007).

34. W. J. Thomas, C. A. Thireault, J. A. Kimbrel, J. H. Chang, Recombineering and stable integration of the Pseudomonas syringae pv. syringae 61 hrp/hrc cluster into the genome of the soil bacterium Pseudomonas fluorescens Pf0-1. The Plant Journal 60, 919–928 (2009).

35. K. Tsuda, I. E. Somssich, Transcriptional networks in plant immunity. New Phytologist 206, 932–947 (2015).

36. D. W. K. Ng, J. K. Abeysinghe, M. Kamali, Regulating the Regulators: The Control of Transcription Factors in Plant Defense Signaling. Int J Mol Sci 19, 3737 (2018).

37. F. Jacob et al., A dominant-interfering camta3 mutation compromises primary transcriptional outputs mediated by both cell surface and intracellular immune receptors in Arabidopsis thaliana. New Phytol 217, 1667–1680 (2018).

38. K. Zhai et al., RRM Transcription Factors Interact with NLRs and Regulate Broad-Spectrum Blast Resistance in Rice. Molecular Cell 74, 996–1009.e1007 (2019).

39. P. D. Townsend et al., The intracellular immune receptor Rx1 regulates the DNA-binding activity of a Golden2-like transcription factor. Journal of Biological Chemistry 293, 3218–3233 (2018).

40. F. Xu et al., NLR-Associating Transcription Factor bHLH84 and Its Paralogs Function Redundantly in Plant Immunity. PLOS Pathogens 10, e1004312 (2014).

41. C. Chang et al., Barley MLA immune receptors directly interfere with antagonistically acting transcription factors to initiate disease resistance signaling. Plant Cell 25, 1158–1173 (2013).

42. X. Liu, H. Inoue, N. Hayashi, C.-J. Jiang, H. Takatsuji, CC-NBS-LRR-Type R Proteins for Rice Blast Commonly Interact with Specific WRKY Transcription Factors. Plant Molecular Biology Reporter 34, 533–537 (2016).

43. X. Chen, C. Li, H. Wang, Z. Guo, WRKY transcription factors: evolution, binding, and action. Phytopathology Research 1, 13 (2019).

44. S. H. Wani, S. Anand, B. Singh, A. Bohra, R. Joshi, WRKY transcription factors and plant defense responses: latest discoveries and future prospects. Plant Cell Reports 40, 1071–1085 (2021).

45. J. Jiang et al., WRKY transcription factors in plant responses to stresses. Journal of Integrative Plant Biology 59, 86–101 (2017).

46. K. Yamasaki et al., Structural basis for sequence-specific DNA recognition by an Arabidopsis WRKY transcription factor. J Biol Chem 287, 7683–7691 (2012).

47. Y.-p. Xu, H. Xu, B. Wang, X.-D. Su, Crystal structures of N-terminal WRKY transcription factors and DNA complexes. Protein & Cell 11, 208–213 (2020).

48. I. Ciolkowski, D. Wanke, R. P. Birkenbihl, I. E. Somssich, Studies on DNA-binding selectivity of WRKY transcription factors lend structural clues into WRKY-domain function. Plant Mol Biol 68, 81–92 (2008).

49. T. Eulgem, P. J. Rushton, S. Robatzek, I. E. Somssich, The WRKY superfamily of plant transcription factors. Trends in Plant Science 5, 199–206 (2000).

50. P. F. Sarris, V. Cevik, G. Dagdas, J. D. G. Jones, K. V. Krasileva, Comparative analysis of plant immune receptor architectures uncovers host proteins likely targeted by pathogens. BMC Biology 14, 8 (2016).

51. Y. Xing et al., Bacterial effector targeting of a plant iron sensor facilitates iron acquisition and pathogen colonization. The Plant Cell 10.1093/plcell/koab075 (2021).

52. J. C. De la Concepcion et al., Polymorphic residues in rice NLRs expand binding and response to effectors of the blast pathogen. Nat Plants 4, 576–585 (2018).

53. J. C. De la Concepcion et al., Protein engineering expands the effector recognition profile of a rice NLR immune receptor. eLife 8, e47713 (2019).

54. J. H. R. Maidment et al., Multiple variants of the fungal effector AVR-Pik bind the HMA domain of the rice protein OsHIPP19, providing a foundation to engineer plant defence. J Biol Chem 296, 100371 (2021).

55. R. Martin et al., Structure of the activated ROQ1 resistosome directly recognizing the pathogen effector XopQ. Science 370, eabd9993 (2020).

56. S. Ma et al., Direct pathogen-induced assembly of an NLR immune receptor complex to form a holoenzyme. Science 370 (2020).

57. J. Wang et al., Ligand-triggered allosteric ADP release primes a plant NLR complex. Science 364, eaav5868 (2019).

58. J. Wang et al., Reconstitution and structure of a plant NLR resistosome conferring immunity. Science 364, eaav5870 (2019).

59. S. Cesari et al., Design of a new effector recognition specificity in a plant NLR immune receptor by molecular engineering of its integrated decoy domain. bioRxiv 10.1101/2021.04.24.441256, 2021.2004.2024.441256 (2021).

60. Á. Piñeiro et al., AFFINImeter: A software to analyze molecular recognition processes from experimental data. Analytical Biochemistry 577, 117–134 (2019).

## References

1. L. E. Bird, High throughput construction and small scale expression screening of multi-tag vectors in Escherichia coli. Methods 55, 29–37 (2011).

2. L. Potterton et al., CCP4i2: the new graphical user interface to the CCP4 program suite. Acta Crystallographica Section D 74, 68–84 (2018).

3. A. J. McCoy et al., Phaser crystallographic software. Journal of Applied Crystallography 40, 658–674 (2007).

4. P. Emsley, K. Cowtan, Coot: model-building tools for molecular graphics. Acta Crystallogr D Biol Crystallogr 60, 2126–2132 (2004).

5. T. I. Croll, ISOLDE: a physically realistic environment for model building into low-resolution electron-density maps. Acta Crystallogr D Struct Biol 74, 519–530 (2018).

6. G. N. Murshudov et al., REFMAC5 for the refinement of macromolecular crystal structures. Acta Crystallographica Section D 67, 355–367 (2011).

7. C. J. Williams et al., MolProbity: More and better reference data for improved all-atom structure validation. Protein Sci 27, 293–315 (2018).

8. G. Battle (2016) PDBePISA: Identifying and interpreting the likely biological assemblies of a protein structure.

9. E. F. Pettersen et al., UCSF ChimeraX: Structure visualization for researchers, educators, and developers. Protein Sci 30, 70–82 (2021).

10. H. Wickham, ggplot2-Elegant Graphics for Data Analysis. Springer International Publishing. *Cham*, Switzerland (2016).

11. Á. Piñeiro et al., AFFINImeter: A software to analyze molecular recognition processes from experimental data. Analytical Biochemistry 577, 117–134 (2019).

12. Panagiotis F. Sarris et al., A Plant Immune Receptor Detects Pathogen Effectors that Target WRKY Transcription Factors. Cell 161, 1089–1100 (2015).

13. Y. Ma et al., Distinct modes of derepression of an <em>Arabidopsis</em> immune receptor complex by two different bacterial effectors. Proceedings of the National Academy of Sciences 115, 10218 (2018).

14. K. H. Sohn, R. K. Hughes, S. J. Piquerez, J. D. G. Jones, M. J. Banfield, Distinct regions of the <em>Pseudomonas syringae</em> coiled-coil effector AvrRps4 are required for activation of immunity. Proceedings of the National Academy of Sciences 109, 16371 (2012).

15. H. Ashkenazy et al., ConSurf 2016: an improved methodology to estimate and visualize evolutionary conservation in macromolecules. Nucleic Acids Research 44, W344–W350 (2016).

